# ASC proneural transcription factors mediate the timely initiation of the neural program during neuroectodermal to neuroblast transition ensuring progeny fidelity

**DOI:** 10.1101/2021.11.02.466869

**Authors:** Vasiliki Theodorou, Aikaterini Stefanaki, Minas Drakos, Dafne Triantafyllou, Christos Delidakis

## Abstract

**Background:** ASC/ASCL proneural transcription factors are oncogenic and exhibit impressive reprogramming and pioneer activities. In both Drosophila and mammals, these factors are central in the early specification of the neural fate, where they act in opposition to Notch signalling. However, the role of ASC on the chromatin during CNS neural stem cells birth remains elusive.

**Results:** We investigated the chromatin changes accompanying neural commitment using an integrative genetics and genomics methodology. We found that ASC factors bind equally strongly to two distinct classes of cis-regulatory elements: open regions remodeled earlier during maternal to zygotic transition by Zelda and Zelda-independent, less accessible regions. Both classes cis-elements exhibit enhanced chromatin accessibility during neural specification and correlate with transcriptional regulation of genes involved in many biological processes necessary for neuroblast function. We identified an ASC-Notch regulated TF network that most likely act as the prime regulators of neuroblast function. Using a cohort of ASC target genes, we report that ASC null neuroblasts are defectively specified, remaining initially stalled, lacking expression of many proneural targets and unable to divide. When they eventually start proliferating, they produce compromised progeny. Generation of lacZ reporter lines driven by proneural-bound elements display enhancer activity within neuroblasts and proneural dependency. Therefore, the partial neuroblast identity seen in the absence of ASC genes is driven by other, proneural-independent, cis-elements. Neuroblast impairment and the late differentiation defects of ASC mutants are corrected by ectodermal induction of individual ASC genes but not by individual members of the TF network downstream of ASC. However, in wild type embryos induction of individual members of this network induces CNS hyperplasia, suggesting that they synergize with the activating function of ASC to establish the chromatin dynamics that promote neural specification.

**Conclusion:** ASC factors bind a large number of enhancers to orchestrate the timely activation of the neural chromatin program during neuroectodermal to neuroblast transition. This early chromatin remodeling is crucial for both neuroblast homeostasis as well as future progeny fidelity.

## BACKGROUND

The Drosophila genome exhibits complex and dynamic developmental chromatin and transcriptional patterns [1-6]. Due to its compact size enhancer elements are tightly spaced and utilized by many, ubiquitous and tissue specific transcription factors (TF) [5, 7-11]. For any given cell-type, specific activators turn on the relevant transcriptional program; while in parallel repressors suppress transcription of genes related to other lineages or temporally inappropriate states, ensuring proper differentiation and maturation [12, 13].

The achaete-scute complex locus (ASC) encodes four paralogous proneural bHLH transcription factors, Achaete (Ac), Scute (Sc), Lethal of scute [L(1)sc] and Asense (Ase), which regulate central (CNS) and peripheral (PNS) nervous system development [14, 15]. They exhibit high evolutionary conservation to mammalian *ASCLs* in both sequence and proneural function [16-21]. Although prominent in neurogenesis, they also regulate progenitor cell specification and function in tissues of endodermal and mesodermal origin [22, 23]. In humans, various studies highlight their oncogenic involvement in malignancies from different germ layers [24]. Examples include small cell lung carcinomas [25], prostate tumors [26], medullary thyroid cancers [27], gastroenteropancreatic tumors [28], gliomas, grade II and grade III astrocytomas and a subset of glioblastoma multiforme [29-33]. Also, their strong reprogramming and pioneer factor abilities [33-37] attest to their transcriptional activating potency.

Within the insects, two ancestral ASC-like proneural factors have been characterized, ASH (Achaete and Scute homologue) and Asense (Ase) [38, 39]. In many insect clades *ASH* genes have duplicated, whereas *ase* has remained as single-copy. Drosophilids three *ASH* genes, *ac, sc* and *l(1)sc*, exhibit a considerable degree of functional redundancy [40, 41]. In the early embryonic neuroectoderm (NE), the naïve CNS primordium, global patterning cues initiate the expression of the three ASH genes in patches of cells [42, 43]. Within these proneural clusters, cells are at a cell fate crossroad, become a neural stem cell, “neuroblast” (NB), and delaminate from the neuroepithelium or remain neuroectodermal and eventually take on the epidermal fate [44, 45]. This cell fate decision is controlled by a finely tuned interplay between ASH proneurals and Notch signalling, mostly through its E(spl)s effectors [14, 46]. Newly born neuroblasts start expressing the fourth paralogue, Ase, and other stem cell markers, and divide asymmetrically to produce ganglion mother cells (GMC), which divide once to produce differentiated neurons and glia. Unlike PNS primordia, where activity of proneural genes is required for precursor specification [15], in ASC-deficient embryos most CNS neuroblasts delaminate, albeit at approximately 25% smaller numbers [47]. These ASC mutant NBs have restricted progeny and often die after stage 11 through a wave of apoptosis. It remains largely unknown how ASC proneurals contribute to CNS neuroblast birth and function at the chromatin level.

Here, we have followed up on early seminal genetic work and addressed this biological process from a genomics point of view and present novel insights regarding the chromatin changes that accompany CNS neural stem cell birth in terms of global proneural binding, active histone mark deposition and transcriptional profiles. Combining these datasets revealed a putative TF-network of proneural target genes, which are likely to comprise the forefront arsenal ensuring neuroblast functionality. Notably, ASC mutant neuroblasts undergo NE to NB transition poorly, remaining in a ‘stalled state’ characterized by lack of timely expression of many proneural targets and, importantly, without dividing. Eventually, they overcome this arrest but cannot sufficiently sustain stem cell competence, evident by the depleted glia and neuronal population resulting in a highly hypoplastic nerve cord. Therefore, ASH proneurals appear to be largely dispensable for the NB delamination process, but are required for timely initiation of the neural stem cell program.

## RESULTS

### Genome-wide mapping of ASH proneural binding during NB specification

To address the role of the ASH proneural factors, we screened a number of Gal4 lines for embryonic neuroectodermal expression and selected bib-Gal4 to express myc-tagged variants of Sc and L(1)sc for genome-wide binding and transcriptome studies. bib-Gal4 is active in the procephalic and ventral neuroectoderm from stage 8 onwards and by stage 16 GFP is detected in the ventral nerve cord (VNC) and the mature epidermis (Fig. 1A, Supplemental Fig. S1). During NB delamination, we detected weak signal in the NBs (Supplemental Fig. S1B), indicating GFP perdurance rather than active GAL4 expression. bib-Gal4 overexpression of a wt Sc did not influence NB specification (not shown). However, induction of scAPAA, a stabilized variant [48], led to a variable, moderate increase in Dpn positive neuroblasts and Pros positive GMCs progeny (Fig. 1B, middle panel). This subtle increase in the NB/GMC population led to mild late-stage CNS hyperplasia (Supplemental Fig. S1C) with varying penetrance and reduced embryonic hatching rate (not shown). On the other hand, overexpression of an extracellular domain deletion of Notch (UAS-NΔecd, abbreviated U-ΝΔE), mimicking Notch activation [49] exhibited reduced number of delaminated neuroblasts (Fig. 1B, bottom panel), severe CNS hypoplasia (Supplemental Fig. S1C-D) and complete embryonic lethality (not shown). These phenotypes agree with the conventional model of mutual proneural - Notch antagonism in NB specification, rendering bib-Gal4 an appropriate driver to monitor the chromatin shifts during NB transition (Fig. 1C).

**Fig. 1.**
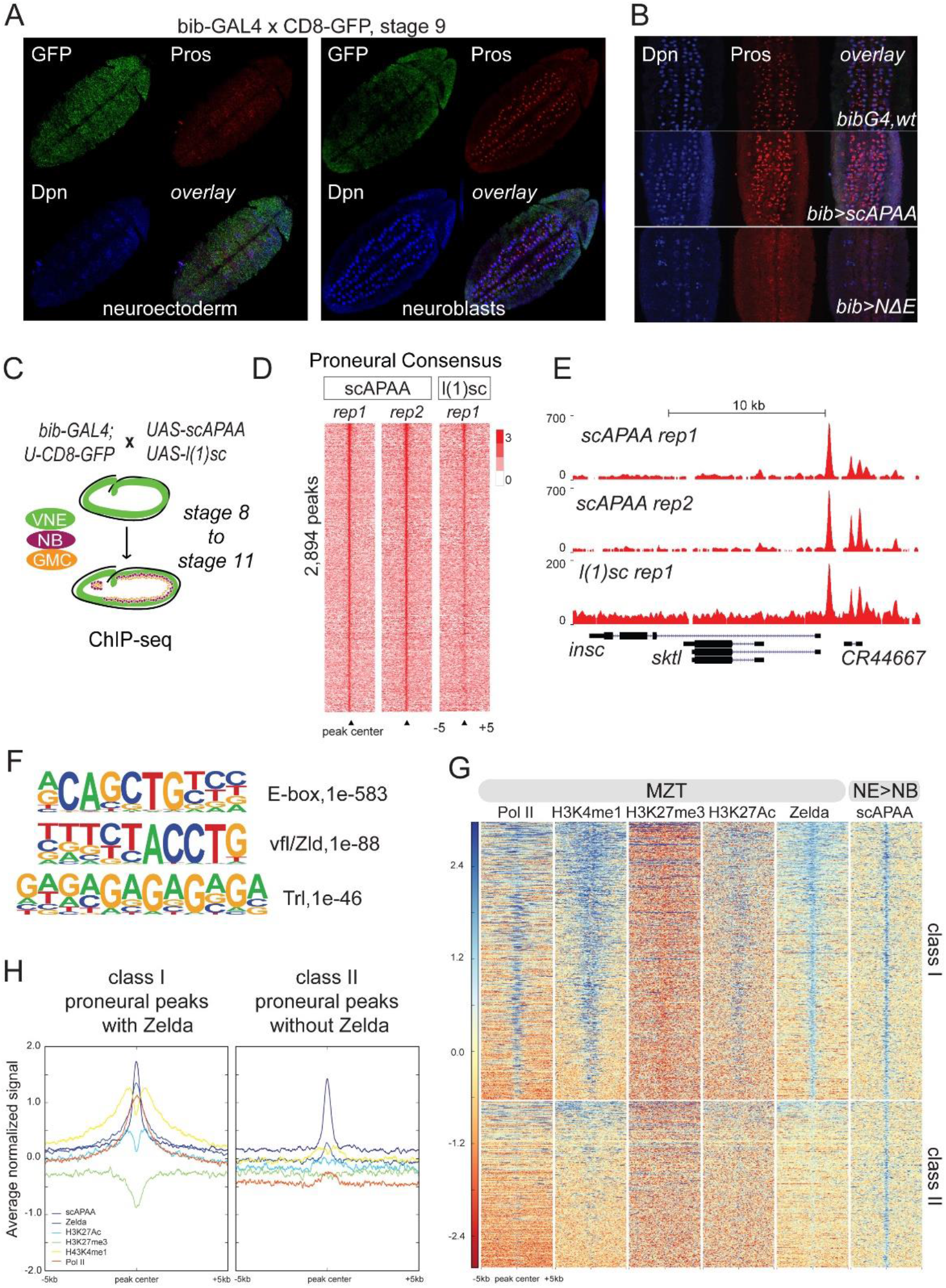
Genome-wide mapping of proneural binding in Drosophila neuroectoderm during neuroblast specification. A) Stage 9 bib-GAL4 embryo shows GAL4 activity in the cephalic and ventral neuroectoderm. B) Close ups in the neuroblast field in stage 9 embryos of the genotypes shown. C) Strategy of staged embryos used as input material to generate the ChIPseq datasets. D) Heatmaps of ChIPseq normalized signal over input centered on the proneural consensus peaks. E) Genomic snapshot at the insc gene. F) De novo motif analysis of the proneural consensus. G) Heatmaps of proneural, Zelda binding, histone marks and poised PolII ChIP-seq signal centered on proneural binding events, grouped in two categories: Class I occupied by Zelda earlier during MZT and Class II, Zelda-independent. H) Average of normalized ChIP-seq signal from heatmaps in G.

We focused on stage 8-mid 11 encompassing almost the entire duration of neuroblast segregation and performed three ChIP-sequencing experiments, two against scAPAA and one against L(1)sc (Fig. 1C). A Venn diagram of called binding events among the three replicates, as well as the signal intensity heatmaps (Supplemental Fig. S1E), show that ScAPAA and L(1)sc bind many genomic loci commonly. We derived a consensus of the two ScAPAA replicates (Supplemental Methods), resulting in 2,894 peaks (Supplemental Table S1). At the level of called peaks, 55% of this strict ScAPAA consensus was also bound by L(1)sc (not shown), possibly due to the overall weaker signal in the l(1)sc library (Fig. 1D).

An example of common proneural binding is shown for the *insc* locus (Fig. 1E). We will refer to this strict, confident consensus of the two ScAPAA replicates as the ‘proneural binding consensus’ for the rest of the paper. This proneural consensus showed 27% overlap with Ac modEncode binding [50] and 12% with the Ase-DamID data [51] (Supplemental Fig. S1F). The limited overlap of ASH proneurals with Ase possibly reflects their expression pattern, since Ase is expressed solely in the delaminated NBs. De novo motif analysis revealed fine differences in the E-box motif for each proneural TF (Supplemental Fig. S1F), highlighting their unique binding preferences beyond their functional redundancy. In addition, we investigated the binding co-occupancy with Daughterless (Da), a well-described proneural partner [52] and E(spl)m8, a neuroectodermal specific Notch induced E(spl) repressor that counteracts proneural/Da function, from modENCODE (Supplemental Fig. S1G). These global comparisons showed a 15% overlap of proneural consensus with Da and 31 % with E(spl)m8, while Da exhibited a much higher, 84% overlap with E(spl)m8 binding events. This raises the possibility that proneurals bind mostly independently of Da and that E(spl)m8 recruitment is channelled through Da rather than proneural factors.

### Proneurals bind developmental DHS regions

Next, we evaluated the genomic distribution of the proneural binding consensus events and found high enrichments in upstream regions (Supplemental Fig. S1H), similar to mammalian Ascl [19] suggestive of an evolutionary conserved positioning of proneural binding motifs close to gene start sites. De novo motif analysis revealed E-boxes as the primary motif identified in 73% of the proneural peak consensus, followed by the Vfl/Zelda and Trl motifs (Fig. 1F). Zelda is the pioneer factor that establishes global chromatin organization during the maternal-to-zygotic transition (MZT) [53-59], which peaks at nuclear cycle 14 (NC14) of stage 5, shortly before ASH expression in the neuroectoderm. Zelda binding together with profiles of various histone modification marks and extensive stalled PolII binding [60-62] has revealed a dynamic chromatin reorganization in preparation for zygotic transcription. We thus overlapped our proneural consensus with stage 5 Zld binding events [56] and found a 62% overlap (Supplemental Fig. S2A), suggesting that at these regions Zelda precedes proneural binding temporally. We used the two classes of proneural bound regions (class I with Zelda, class II without Zelda) to investigate the chromatin landscape patterns prior to proneural binding. Based on the patterns of H3K4me1 and H3K27Ac, positively associated with chromatin accessibility, the lack of the repressive H3K27me3, and the PolII signal it appears that prior to proneural binding class I target regions were nucleosome remodeled and more accessible whereas class II sites were less accessible. Subsequently, during NB specification proneurals appear to bind these loci equally strong (Fig. 1G-H). These two classes of cis-elements exhibited differences in motif enrichment analysis suggesting possible differential TF recruitment (Supplemental Table S2). Also, class II elements were less frequently located within a 5kb window upstream from the TSS (Supplemental Fig. S2B) suggesting that they constitute long-range, tissue-specific enhancers (Reddington 2020).

Since regulatory elements correlate with DNAse Hypersensitivity Sites (DHS) [8, 11] we investigated proneural binding occurrence within stage specific DHS and found striking overlaps (Supplemental Fig. S2C-D). Notably, 89% of proneural binding events were within DHS from all stages, with higher overlaps in stages 9-11 in agreement with proneural activity during NB specification. The vast majority, 98%, of class I proneural events were within DHS (Supplemental Fig. S2C), while class II exhibited a smaller overlap at 74% (Supplemental Fig. S2C). Importantly, Class I elements were open from st5 onwards, whereas Zelda-independent Class II elements were more dynamic, becoming more accessible as embryos progress from st5 to st11, perhaps as a result of proneural pioneer activity in preparation for the neural-specific transcriptional program.

### Proneurals target a plethora of genes necessary for proper NB homeostasis

We then assigned the proneural consensus binding events to 1,983 genes and used the Flymine tool [63] for downstream mining (Supplemental Table S3). Gene Ontology analysis (Fig. 2A) showed high enrichments for nervous system development and DNA-binding transcription factors. 53 members of the Homeobox-like domain superfamily, 69 Zinc finger C2H2-type and 21 Helix-loop-helix DNA-binding domain superfamily genes were amongst the proneural targets, suggesting proneural regulation of a broad network of transcription factors.

**Fig. 2.**
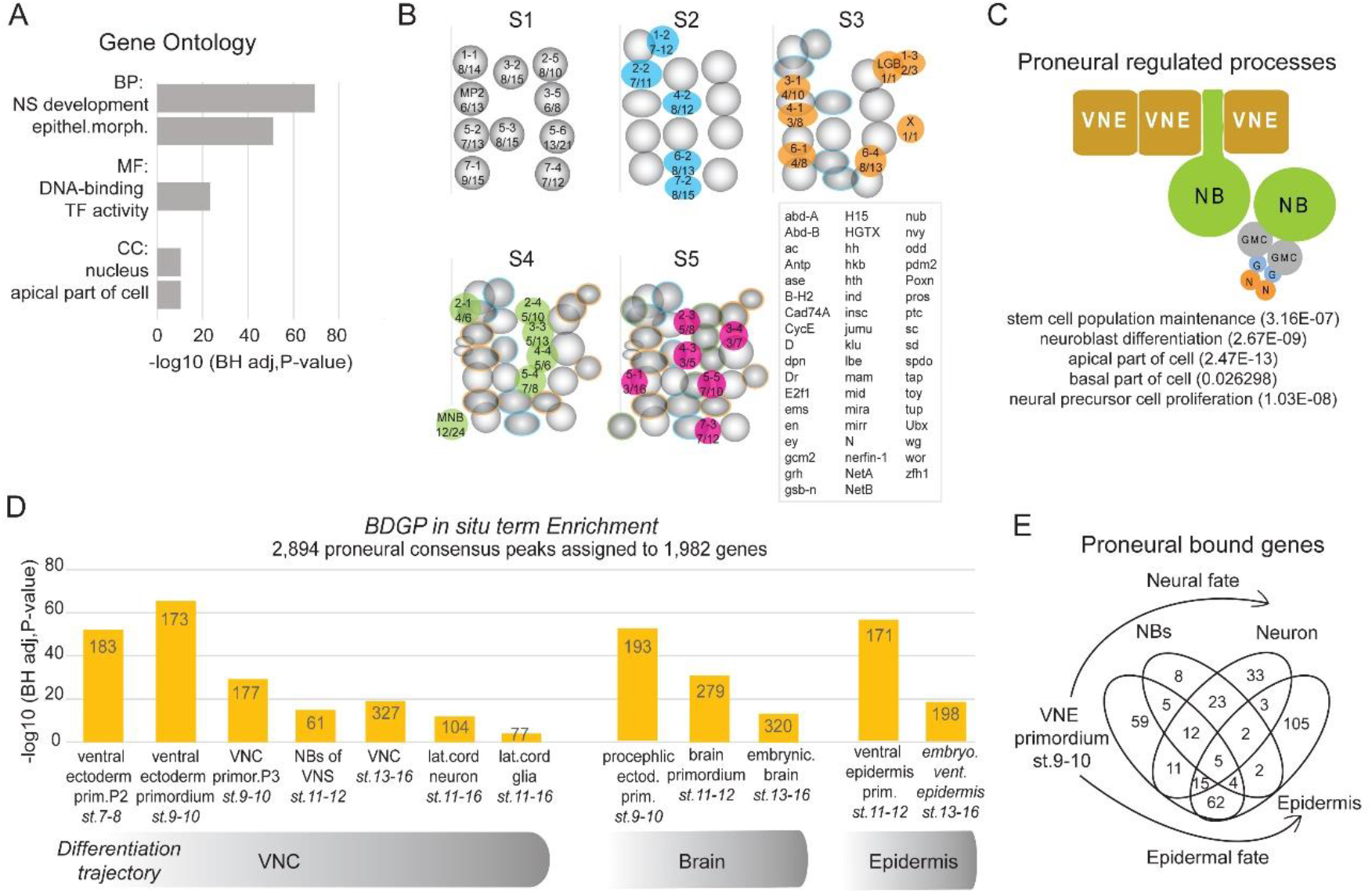
Proneurals target many genes and pathways that convey neuroblast homeostasis. A) Gene Ontology analysis of proneural targeted genes, Biological Processes (BP), Molecular Function (MF) and Cellular Compartment (CC). B) Overlap of Flybase neuroblast genes with proneural targets shown in the 5 consecutive waves of NB specification S1-S5. Numbers under the neuroblast IDs represent the number of proneural targets over the total Flybase NB specific genes. Boxed inset lists the sum of the proneural bound neuroblast markers. C) Proneurals regulate a holistic neuroblast program. A schematic summary of selected terms. D) BDGP in situ enrichments of proneural target genes. E) A venn diagram of proneural bound genes from the BDGP database in D.

Next, we extracted from Flybase [64] genes associated with each specific neuroblast and found proneural binding in 53 out of 98 neuroblast markers, of all five waves of neuroblast specification (Fig. 2B). Besides genes that presumably provide neuroblast identity (stemness), many different processes are needed for proper NB function: delamination; establishment of cytoplasmic asymmetry, expression and correct segregation of pro-differentiation factors, self-renewal and proliferation through multiple asymmetric divisions and temporal progression of progeny types [14, 65]. Notably, proneural target genes fell in all above-mentioned processes. For instance, the known stem cell identity markers *wor, dpn, scrt, klu*, the temporal genes *hb, Kr, nub, grh* [66], genes encoding myosin contractile machinery important for delamination, like *zip, sqh, Rok* and *Rho1* [67], the cell cycle genes *cycE, E2F1* and *stg* and members of apico-basal polarity organizing Par complex (*baz*), Pins complex (*insc,loco,mud,cno*), the Centrosome organizing center (*ctp, mud*) and the basal compartment (*mira, brat, pros*) [68]. Thus, proneurals appear to regulate besides delamination many biological processes needed for neuroblast homeostasis (Fig. 2C).

In addition, we investigated the expression patterns of the proneural-targeted genes using the BDGP *in situ* RNA database integrated in the Flymine tool (Fig. 2D, Supplemental Table S3). We found that many target genes express in the ventral ectoderm primordium, but also in brain, VNC, midline and sensory primordia at the time of neural specification. We also found binding near genes expressed in later developmental stages, in differentiated cell types such as neurons and glia, also supported by the GO enrichments in neuron differentiation [GO:0030182] and axonogenesis [GO:0007409] (Supplemental Table S3). A venn diagram of proneural-bound genes, expressed in the ventral ectoderm, NB, VNC neurons and epidermis (BDGP), showed common as well as unique genes per cell type (Fig. 2E). Thus, we speculate that besides orchestrating the neuroblast program, during the NE to NB transition, proneurals may remodel chromatin in preparation for more committed differentiation states.

### Proneural binding enhances chromatin acetylation

Next, we asked whether proneural activity affects chromatin organization in terms of enhancer remodeling and transcriptional output. For this reason, we generated four replicated RNA-seq experiments and an H3K27Ac ChIP-seq dataset from staged embryos (Fig. 3A). We restricted the time window for these experiments by 1 hour (stage 8-mid 10) compared with the proneural ChIP-seq datasets, to ensure monitoring the initial process of NE➔NB specification and dilute out possible signal from more differentiated cell types. First, we focused on the proneural peak consensus and found higher H3K27Ac signal in the U-scAPAA embryos, in both class I and class II regions (Fig. 3B). Importantly, class II elements, which at NC14 exhibited overall low accessibility, had undergone nucleosome remodelling by st10 (compare the shapes of averaged signal in NC14 Fig. 1H to Fig. 3B) and exhibited increased H3K27Ac signal in UAS-scAPAA embryos compared to wt or UAS-NΔE. Genomic snapshots at *wor* and *nvy*, two bona fide neuroblast markers [69, 70] are representative examples (Fig. 3C). Along this line, analysis of H3K27Ac mark on st9 DHS sites, revealed increased signal in the proneural-bound DHS regions (left panel) compared to the non-bound DHSs (middle panel) (Fig. 3D and Supplemental Fig. S2E). This indicates that Drosophila proneurals elicit nucleosome remodeling and enhance active chromatin conformation, consistent with the pioneer function of mammalian homologues [36, 37].

**Fig. 3.**
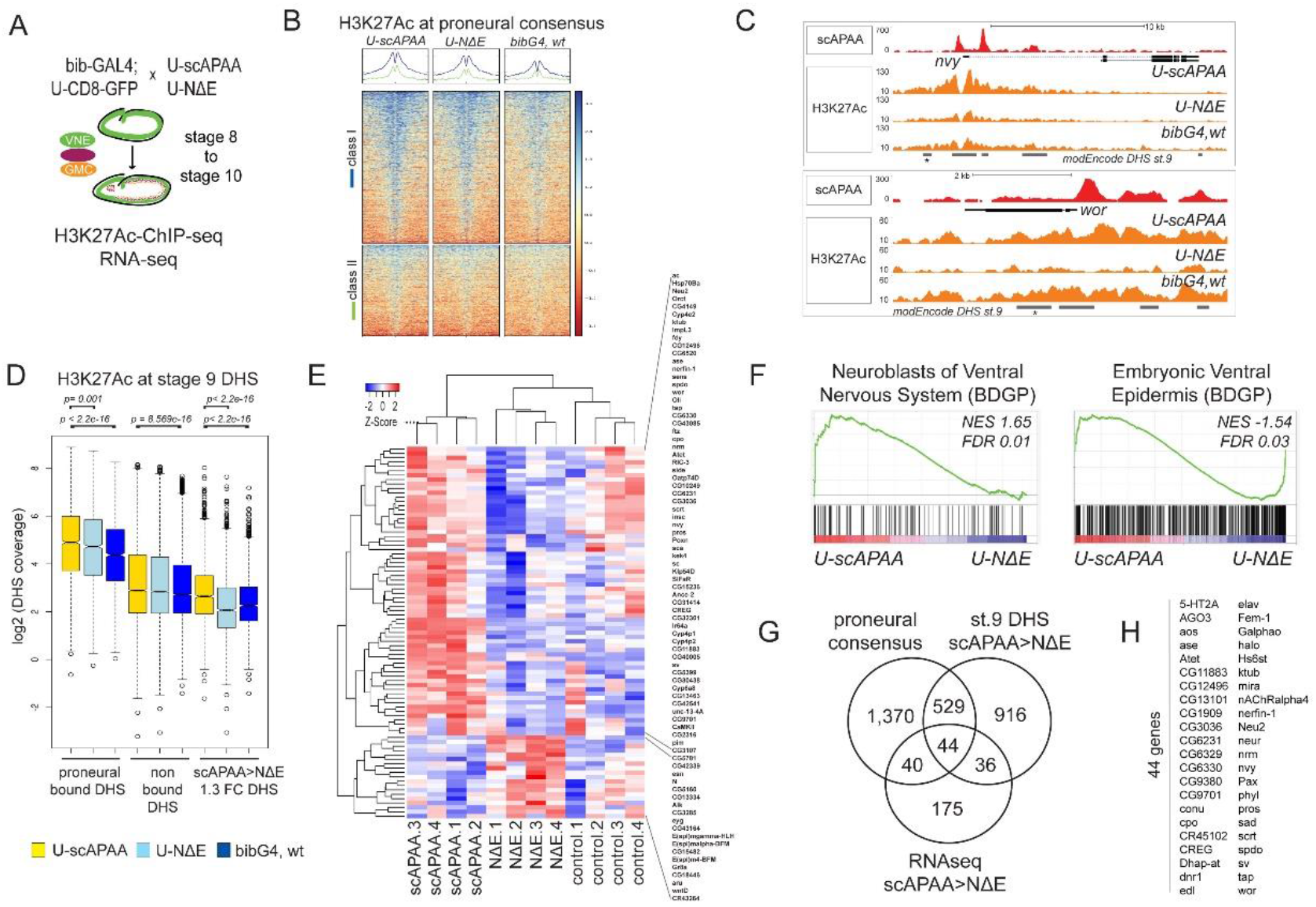
Proneural mediated chromatin changes correlate with transcriptional output during early neurogenesis. A) A schematic representation of the strategy used to generate H3K27Ac ChIP-seq datasets and RNA-seq profiling. B) Heatmaps of H3K27Ac ChIPseq signal centered on Class I and Class II proneural peaks. (C) Genomic snapshots at the nvy and wor loci. * mark DHS st.9 sites without proneural binding that exhibit increase in H3K27Ac signal in U-scAPAA vs. U-NΔE. D) Boxplots of normalized H3K27Ac signal in stage 9 DHS sites from modENCODE. DHS with proneural binding (left), not proneural-bound DHS (middle) and DHS sites that exhibited more than 30% difference in U-scAPAA versus U-NΔE conditions (right). Statistics performed with Wilcoxon rank sum tests. E) Differential Expressed Genes in scAPAA versus NΔE embryos FDR 0.2F) Gene Set Enrichment Analysis (GSEA) of RNAseq data reveal enrichment for neuroblasts and ventral epidermis. G) A venn diagram of genes with proneural binding, the affected DHS sites from E (right) and differential expressed genes (p<0.05) form the RNAseq. H) List of the 44 target genes from the intersection in G.

We subsequently asked which DHSs were most affected in scAPAA vs. NΔE conditions, as a way to monitor the neuroblast versus epidermal cell fate selection during lateral inhibition. 1,889 loci exhibited more than 30% positive difference in active histone deposition in scAPAA versus NΔE overexpressing embryos (Fig. 3D, right panel, and Supplemental Fig. S2E). These genomic sites were near 1,525 genes, enriched in ventral ectoderm and nervous system related genes (not shown), similar to the proneural consensus distributions of Fig. 2D. However, only 16%, (306 sites), of the affected DHSs coincided with proneural binding (not shown). The remaining not-bound DHSs were close to proneural-bound genes, 39% overlap at the gene assignment level, which indicates that proneural binding has broader effects outside its binding element, either as a result of gene transcription or long-range looping interactions (note the non bound DHSs with * at the wor and nvy examples in Fig. 3C). Alternatively, these differentially acetylated DHSs may represent cis-elements regulated by Notch signalling independently of ASH activity.

### Combination of transcriptome and chromatin profiling reveals putative core regulators of neural stem cell function

To identify the transcriptional changes that accompany neural selection, we performed RNAseq expression profiling (Supplemental Table S5). In the differentially expressed genes (DEG) between the U-scAPAA and U-NΔE embryos (FDR<0.2, p<0.0025) (Fig. 3E) there were many neurogenesis related transcription factors. Indeed, the ranked genes clearly mirrored the neural versus epidermal fate specification that proneurals and Notch favor respectively (Fig. 3F). In addition, we found significant enrichments with the class II proneural binding events (Supplemental Fig. S2F) as well as with the affected DHSs in H3K27Ac deposition in scAPAA>NΔE (Supplemental Fig. S2G). These correlations demonstrate that the regulatory elements filtered out from the above integrative genomics analyses are transcriptionally relevant. To expand on this observation, we overlapped (a) genes with higher RNA expression in U-scAPAA versus U-NΔE embryos (with the significance threshold relaxed to p<0.05), (b) genes that exhibited proneural binding and (c) genes with differential H3K27Ac in nearby DHSs (Fig. 3G). We found that 40% of DEGs were associated with the one or/and the other dataset. In Fig. 3H we present the 44 genes from the intersection of the three. In this high-confidence gene set we find *ase, nerfin-1, sv, tap, pros, sens, scrt, wor* and *nvy*, known to act in the CNS, PNS and midline. Thus, this TF network regulated by proneural and Notch interplay could be the initial battery of factors required to sustain neural precursor functionality.

### ASC mutant neuroblasts are temporarily stalled and devoid of stem cell identity markers

ASC null (Df(1)scB57) embryos show a reduced number of delaminated NBs and a drastic reduction of mature neurons [47]. However, it has not been documented in detail how these mutant NBs behave. For this purpose, we selected 13 TFs (Dpn, Klu, Wor, Sna, Esg, Scrt, Nvy, Pros, Hb, Kr, Nerfin-1, Tap, Oli), whose genes exhibited proneural regulation in our genomic analyses, and examined their expression in wt vs ASC null embryos. In wt embryos all 13 display NB expression to some extent. A summary of their genomic features and expression patterns is in Supplemental Table S6.

Unexpectedly, in the ASC deletion we observed that delaminated neuroblasts are temporarily stalled during stages 9 and 10. They do not express the stem cell specific markers Dpn (Fig. 4A), Wor (Figute S3A), Nvy (Fig. 4C), Scrt (Fig. 4E), Klu and Oli (not shown), compared to wt NBs which during this time robustly express all these TFs. In contrast, the expression of Hb (Fig. 4A), Sna, Esg and Kr (not shown), appeared unaffected in the mutant NBs. Significantly, mutant neuroblasts did not proliferate, evident by the lack of Pros positive GMCs (Fig. 4A) and phosphoH3-S10 (pH3) mitotic events (Supplemental Fig. S3B). We used the UAS-FUCCI, a GFP-E2F1 and RFP-CycB dual expressing system, that allows cell cycle monitoring by fusing cell-cycle specific degrons to fluorescent proteins [71]. Consistent with bib-Gal4 activity specifically in the NE, wt NBs showed little or no accumulation of FUCCI signal (Supplemental Fig. S3C). Stalled ASC NBs, however, accumulated both these markers demonstrating a G2/M arrest, suggesting that after delamination they retained the NE-expressed FUCCI signal since they had not divided yet (Supplemental Fig. S3D). These results suggest that ASC deficient neuroblasts undergo NE to NB transition poorly as they do not proliferate, nor initiate expression of the entire neural TF program (Supplemental Fig. S3E).

**Fig. 4.**
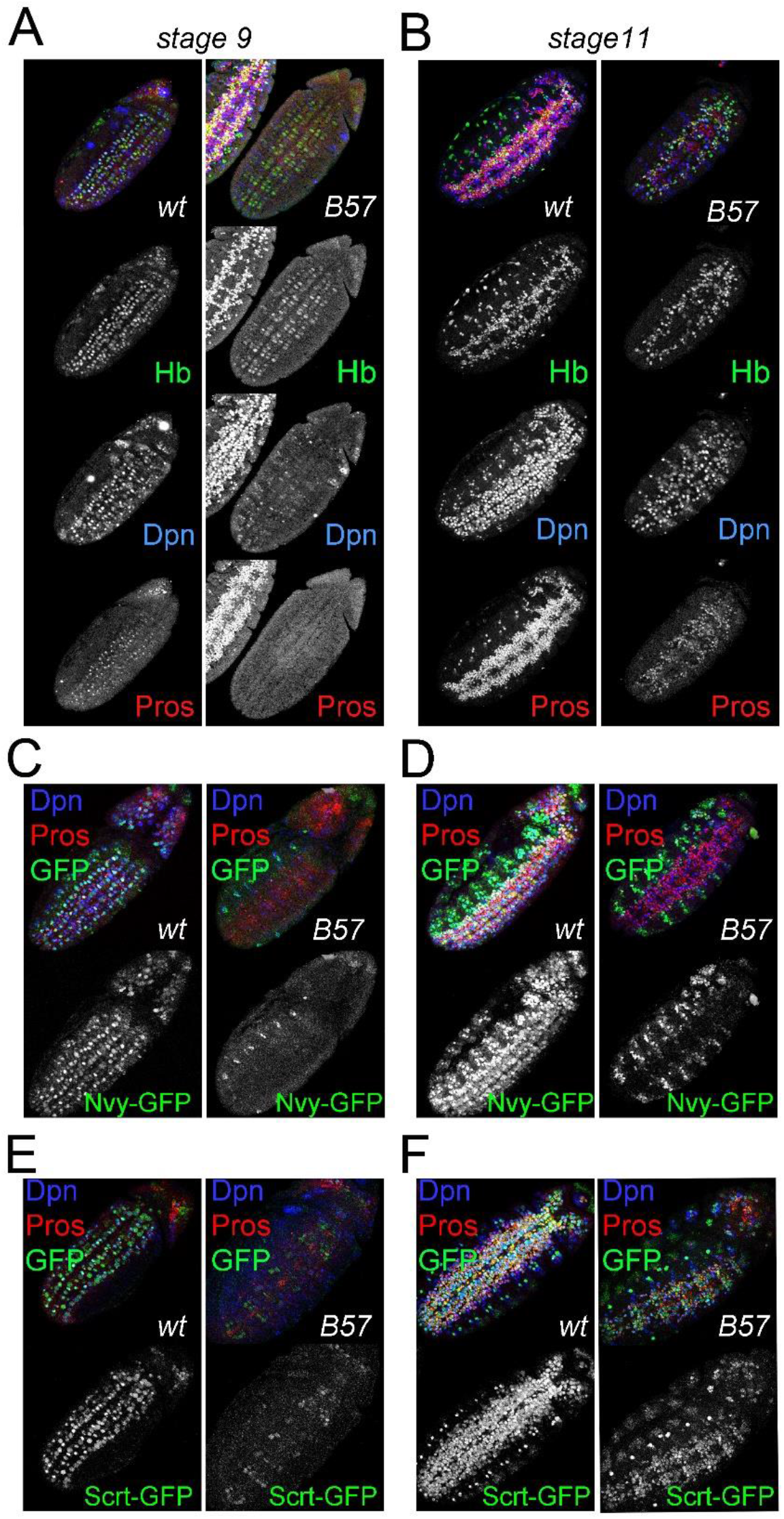
ASC mutant neuroblasts are temporarily stalled and devoid of stem cell identity markers. A) Stage 9 wt neuroblasts (left panels) express Hb and Dpn and have divided to generate Pros positive GMCs. In Df(1)scB57 embryos (right panels) neuroblasts express Hb but not Dpn and have not yet produced GMCs. The weak Dpn signal in the mutant embryo comes from the NE layer above the delaminated NBs. B) In stage 11 mutant neuroblasts have rebounded in Dpn expression and cell divisions to produce GMCs. The sparse Dpn and Pros positive cells outside the broad band of the VNC are PNS precursors, which are also strongly reduced in the ASC mutant. C) Nvy-GFP is absent in mutant neuroblasts during stage 9. Remainig expression comes from more laterally positioned PNS precursors, D) Nvy expression does not rebound in mutant neuroblasts at st 11. E) Scrt is lost or very weak in mutant neuroblasts at st 9. F) Scrt expression rebounds in stage 11 Df(1)scB57 neuroblasts and GMCs.

Despite this early developmental arrest, starting at late stage 10/early 11, we observed a gradual rebound in NB marker expression, accompanied by initiation of NB mitoses. By late stage 11, mutant NBs started expressing Dpn (Fig. 4B), Scrt (Fig. 4F), Oli (Supplemental Fig. S4), Wor and Klu (not shown). The only marker that never rebounded, demonstrating obligate ASC NB regulation, was Nvy (Fig. 4D). Hb (Fig. 4B) and Sna (not shown), not affected at earlier stages, were turned off as usual at this late stage, while Kr continued expressing from earlier stages as normal (not shown). Concomitantly, many GMCs were born, albeit with an aberrant molecular profile. These GMCs expressed Pros (Fig. 4B), Scrt (Fig. 4F), Esg, Hb (subset, Fig. 4B), Kr (subset, not shown), Oli (subset, Supplemental Fig. S4A) and Nerfin-1 (subset, Supplemental Fig. S4B), but not Nvy (Fig. 4D) or Tap, which is mostly expressed in a large subset of GMCs (Supplemental Fig. S4C). Tap eventually turned on in GMCs by stage 13 (not shown). Therefore, the timely expression of neuroblast and GMC markers and proliferation capacity of neural stem cells is ASC dependent.

### ASC mutant neuroblasts are defective and produce impaired progeny

Despite this rebound in mutant NB identity, late embryos are severely hypoplastic. Staining with axonal markers revealed a fragmented nerve cord, a complete lack of the three VNC longitudinal nerve tracts and severe defects in intersegmental/segmental nerves (Supplemental Fig. 5A), see also [47, 72]. Axonogenesis is normally guided by communication cues between neurons and glia from the CNS, PNS and midline [73-78]. Glia play a crucial role both in prefiguring axonal paths and in providing trophic support to neurons. This is evident in glia depleted, gcm mutant embryos [79], where longitudinal nerve tracts also fail to develop. We found a diminished glia population in late ASC embryos. This was more evident in the abdominal segments, by an at least 70% reduction in Repo positive glia (Supplemental Fig. 5B). Specifically, the two characteristic continuous columns of longitudinal glia lining the dorsal side of the developing nerve cord from st13 onwards were depleted. Their longitudinal glioblast progenitor (LGB), however, was present in many hemisegments earlier (st.10/11) (not shown); suggesting that in ASC mutants the born LGB is defectively programmed.

Besides gliogenesis, we next assessed the ability of mutant NBs to generate pioneer neurons that, together with the depleted glia, would explain the lack of longitudinal nerves. It is already known that two pioneer sibling neurons, dMP2 and vMP2 (progeny of MP2, an S1 wave NB), are absent or mis-specified in ASC mutants [80, 81]. We used Eve staining, to identify the aCC/ pCC sibling pioneer neurons (progeny of S1 NB1-1), as well as the U-neurons (S1: NB7-1), the EL-neurons (S4: NB3-3) and the RP2 motor neuron (S2: NB4-2) (Supplemental Fig. 5C). ASC stage 11 embryos have only just started producing GMCs, accordingly no Eve positive neurons were seen (not shown), a time when normally the aCC/pCC pair is formed and expresses Eve and Fas2 [82]. In later stages, we still failed to detect the Eve+ aCC/pCC pioneer neuronal pair (Supplemental Fig. 5C). Only one medial Eve positive, Fas2 negative neuron was observed (not shown), presumably the RP2. In addition, we observed reduced numbers of EL neurons and extremely rare U neurons (Supplemental Fig. 5C).

Since the four pioneer neurons (aCC/pCC and dMP2/vMP2) and the longitudinal glia are born from precursors specified at the early S1-S3 waves of neurogenesis, we wondered whether ASC mutants also exhibit defects in neurons/glia born from later NB waves during late st10-11, a time when mutant NB activity has rebounded. eagle-lacZ is a marker of four NBs and their progeny, three of which arise during S4-S5 (S3:NB6-4, S4:NB2-4, and NB3-3 and S5:7-3) [83]. We observed that in mutant embryos these NBs delaminate and are present in most neuromeres (Supplemental Fig. S6A-B), however, their progeny is variably depleted (Supplemental Fig. S6C-D) and their axonal projections deformed, accompanying an anterior to posterior commissure (AC-PC) collapse (Supplemental Fig. S6E). Collectively, these observations suggest that ASC deficient NBs, both from early and late phases of specification, have an inherently defective program and cannot sustain correct progeny differentiation.

### Ase can substitute for the ASH genes to initiate the neural program in the neuroectoderm

We next investigated whether any of the downstream proneural targets revealed by our genomic experiments would be able to rescue the neurogenesis defects of the ASC deficiency, if transgenically provided using the neuroectodermal driver bib-Gal4 (Fig. 5). We tested UAS-scrt, UAS-wor, UAS-dpn and UAS-Oli, four of the proneural targets that showed a delayed onset of expression in the absence of the ASC. None of these was able to rescue NB stalling at st9. We observed a detectably earlier rebound of NB activity at early st10, evident by the earlier Dpn expression and the emergence of Pros+ GMCs (Fig. 5B-E, top panels). Nonetheless, this slight NB rescue was not able to improve the severe late hypoplastic phenotype (Fig. 5B-E, bottom row), suggesting that these factors are not capable of activating the full neurogenic program in the absence of ASC genes. In contrast, induction of UAS-scAPAA or UAS-ase led to a vast improvement in the delamination defect and the timely activation of NBs (Fig. 5F-G, top), which now started dividing normally at st9. At later stages, the VNC was almost complete with only minor constrictions (Fig. 5F-G, bottom).

**Fig. 5.**
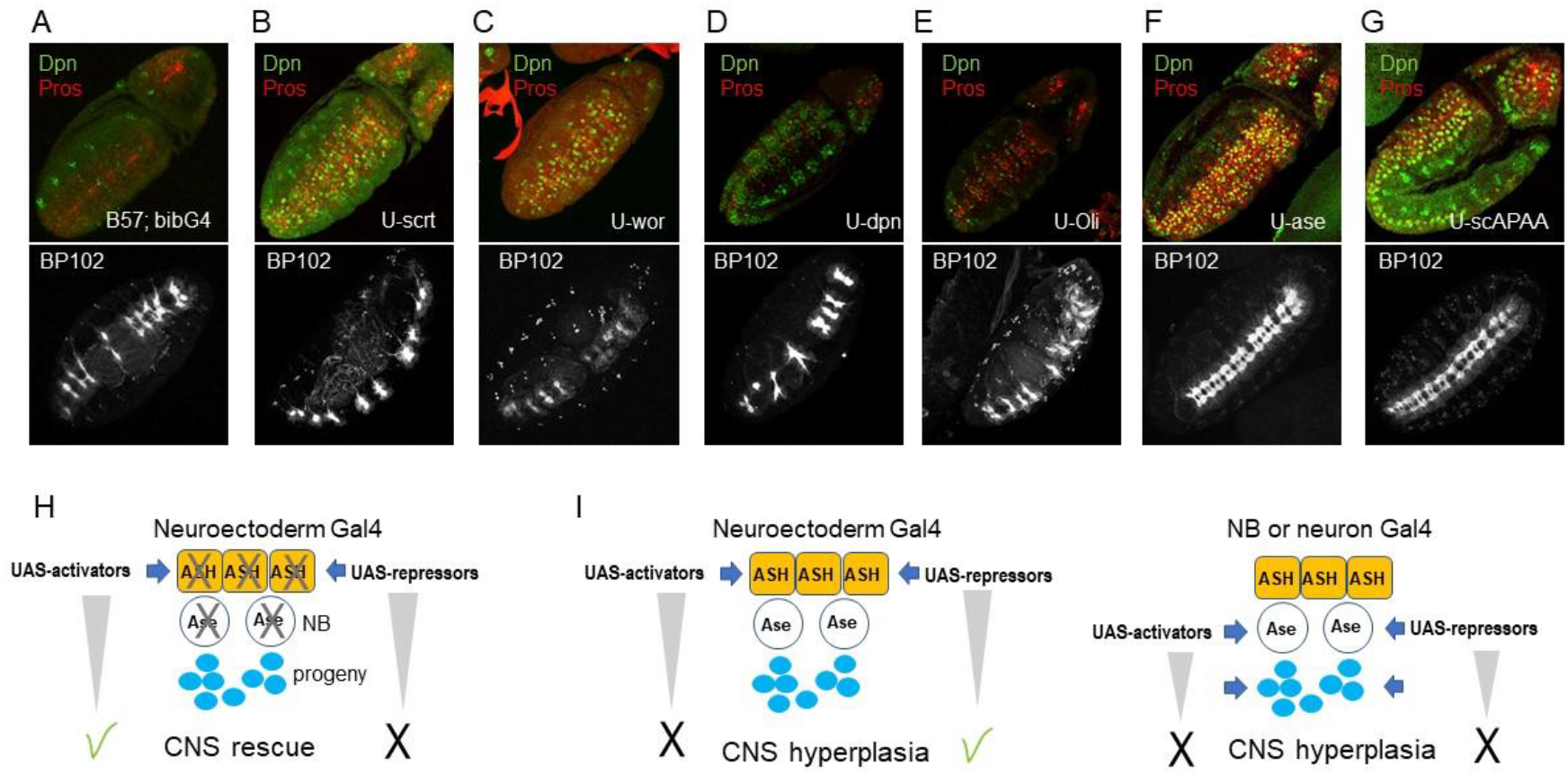
ASC loss is hard to compensate. Early and late rescue phenotypes of neuroectodermally (bibGal4) induced proneural targets in the Df(1)scB57 background. Early embryos (top row) stained with Dpn and Pros, late embryos (bottom row) stained with the axonal marker BP102. A) Df(1)scB57; bibGal4 with no UAS transgene. B-G) as in A, plus B) UAS-scrt C) UAS-wor D) UAS-dpn E) UAS-Oli F) UAS-ase G) UAS-scAPAA. H) Model of ability of selected genes to rescue the Df(1)scB57 neuronal hypoplasia I) Model of ability of selected genes to induce neuronal hyperplasia in the wt background. Activators refer to ASC genes; repressors refer to Snail and Hes family genes. Effect is shown by a check mark; lack of effect by X.

Therefore, re-instating proneural expression in the neuroectoderm can greatly rescue neurogenesis demonstrating that the ASH and Ase proteins have equivalent activities, despite their distinct expression patterns. To clarify this further we used the Df(1)sc19 ASC deficiency (Supplemental Fig. S7), which deletes *ac, sc* and *l(1)sc*, but spares *ase*. In this background, NB stalling was still evident during stage 9 (Supplemental Fig. S7A). Ase itself also exhibited a small delay in expression, however its expression preceded Dpn (Supplemental Fig. S7B) and Pros (not shown), both rebounding a little after Ase expression by early stage 10 (Supplemental Fig. S7C), earlier than in Df(1)scB57. The late CNS hypoplasia was also improved in Df(1)sc19. The population of glia was richer (Supplemental Fig. S7D) and the aCC/pCC pioneer neuron pair was sometimes present (not shown). The VNC had fewer neuromere gaps, as reported by [72], although the wt pattern of three Fas2-bearing longitudinals was never fully restored (Supplemental Fig. S7E). Therefore, the endogenous expression of Ase in the delaminated neuroblasts can greatly improve NB functionality (sc19 vs.B57), but not as efficiently as when we induce it in the neuroectoderm during NB specification (Fig. 5F), suggesting that the neuroblast program at the chromatin level commences during the NE to NB transition.

The foregoing experiments demonstrated that although individual ASC proneurals are sufficient to rescue the CNS defects caused by ASC deletion, none of their other primary targets tested were competent to do so (Fig. 5B-E). However, in the presence of proneural proteins (in wt background), scrt, wor and dpn neuroectodermal overexpression by bib-Gal4 led to significant neural hyperplasia evident at the level of longitudinal connectives and segmental/ intersegmental nerve bundles (Supplemental Fig. S8A). Cuticle preps showed epidermal holes (Supplemental Fig. S8B), suggesting that scrt, wor or dpn NE overexpression tipped the balance in favour of NBs at the expense of epidermis. Although, in the wild type context bib>scAPAA overexpression on its own had a weak effect (Supplemental Fig. S1B-C), coexpression with dpn enhanced the hyperplasia produced by either alone (Supplemental Fig. S8A). Similar enhancement was observed upon co-expressing two proneurals together, scAPAA with l(1)sc (Supplemental Fig. S8A). Notably, VNC hyperplasia was not seen when these genes were induced in the neuroblasts by pros-Gal4 (starts expressing in st11 NBs, GMCs and neurons) (Supplemental Fig. S8C) or in neurons using elav-Gal4 (starts expressing in st13 NBs, GMCs and neurons, not shown). These results suggest that TFs of the Snail (Wor, Scrt) and Hes families (Dpn), most known to act as repressors [84, 85], can enhance the NB-promoting activity of proneural TFs, but have little genuine activating potency to initiate the neural program on their own (Fig. 5H-I, model cartoons). This conclusion is supported by the ectopic neural cells in the wing disk induced by a TF cocktail consisting of a proneural (Ase), a Snail (Wor), as well as two more broadly NE-expresssed TFs (SoxN and Kr) [86].

### Proneural bound cis-elements exhibit enhancer activity and proneural dependency

To investigate the transcriptional activity of the proneural bound elements we generated 10 transgenic lacZ reporter flies. We selected proneural peaks, near *nvy, dpn, scrt, wor* and *tap* genes, whose protein products showed proneural dependency in mutant embryos in our foregoing analysis. We included binding events near *insc* and *brat*, two key neuroblast genes that are implicated in apico-basal polarity and asymmetric cell division [87] and one intronic peak from the *phyl* gene, a known PNS proneural target [88]. Most of these regions coincided with DHS sites and half had Zelda binding during MZT (Supplemental Table S7). All fragments showed enhancer activity in some regions of the developing nervous system, central and/or peripheral and none in non-neural tissues. The wor-KV29 exhibited weak expression and was not studied further. For the remaining lines, we compared the lacZ expression patterns in wildtype and Df(1)scB57 embryos, summarized in Fig. 6A.

**Fig. 6.**
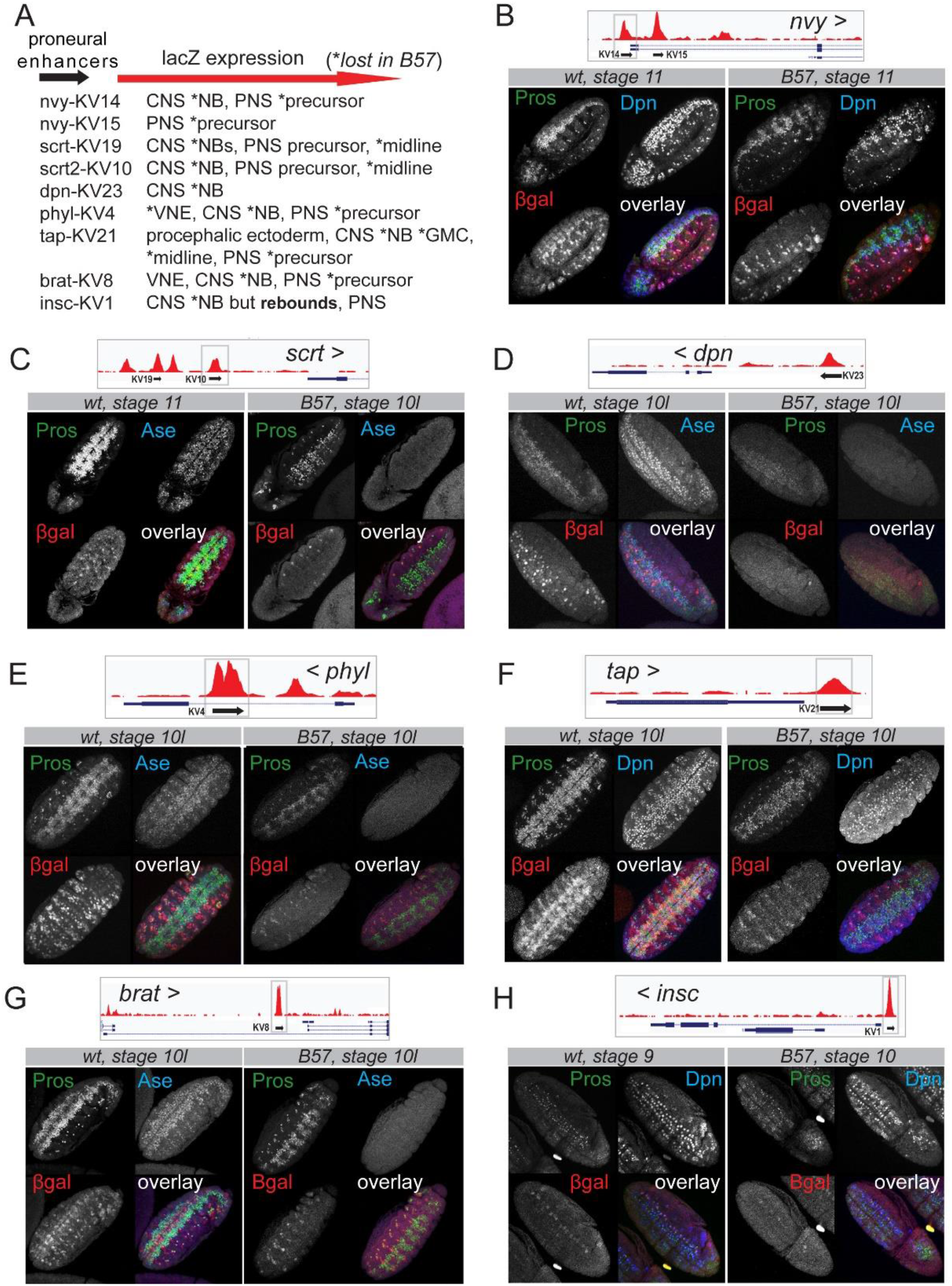
Proneural bound genomic elements exhibit spatiotemporal enhancer activity and proneural dependency. A) Summary of enhancer spatiotemporal expression patterns in wt and Dfsc(1)scB57 (*) embryos. B) Embryos expressing the upstream nvy-KV14 reporter. C) Embryos expressing the upstream scrt-KV10 reporter. D) Embryos expressing the upstream dpn-KV23. E) The intronic phyl-KV4 reporterin stage 10 embryos. F) The 3’ prime tap-KV21 reporter. G) The KV8 reporter proximal to the short brat isoforms H) The proximal to TSS insc-KV1 reporter. In the genomic insets, black arrows indicate the extent and cloning orientation of the genomic elements in the lacZ expressing vectors. The > symbol next to gene names shows the orientation of transcription.

Briefly, the *nvy* enhancers, exhibited different patterns, nvy-KV14 had CNS and PNS expression (Fig. 6B) while nvy-KV15 was PNS exclusive (not shown). In mutant neuroblasts, nvy-KV14 expression was abolished throughout neurogenesis similar to the Nvy protein (Fig. 4C-D). In scratch-KV10-lacz wt embryos, we detected moderate NB and stronger midline signal, which was lost in mutants (Fig. 6C) in contrast to the rebound in scrt-GFP protein (Fig. 4F). dpn-KV23, was expressed in S3 and S4 NB waves and by stage 13 had expanded to cover the whole NB pool (not shown). In the Df(1)sc-B57 mutant KV23 was never activated (Fig. 6D), in contrast to the resumed Dpn protein expression (Fig. 4). The phyl-KV4 enhancer expressed from st9/10 in NBs and some VNE clusters (Fig. 6E). Next, tap-lacZ, exhibited ectodermal, CNS (subset of NBs and GMCs) and PNS expression (Fig. 6F). In early mutant embryos, the NB/GMC expression was lost (Fig. 6F) but we did detect limited expression in GMCs and midline from stage 13-14 onwards (not shown). Similarly, brat-lacZ (Fig. 6G) exhibited broad neuroblast expression in wt embryos but its expression was lost in the mutant background, even after the onset of asymmetric divisions and generation of Pros positive GCM progeny. Lastly, the insc-KV1 enhancer showed extensive NB expression from S1-S2 onwards with an emphasis in the lateral and intermediate rows. It exhibited absence of expression in mutant NBs during the stalling window but did express during the rebounding period (Fig. 6H). Thus, with the sole exception of the *insc* enhancer, the NB-specific activity of *nvy, scrt, dpn, phyl, brat* and *tap* regulatory elements exhibited absolute ASC dependency both during stalling as well as after stem cell activity resumption. This suggests that, at the chromatin level, the delayed NB activation in the absence of proneurals is mediated by cis-elements distinct from those bound by proneural proteins. Unlike NB expression, all enhancers that drove PNS expression displayed activity in the Df(1)scB57 mutant in the ASC-independent sensory organs [89], most likely due to the activity of the *atonal* and *amos*, proneural factors exclusive to PNS primordia [90, 91].

## DISCUSSION

### Chromatin dynamics during embryonic nervous system development

The advent of genomics has revealed that the chromatin of any given cell is a blueprint of past, current and future maturation states both in homeostasis and disease [92-96]. In Drosophila as well, studies of cell fate transitions depict dynamic chromatin shifts during development [2-5, 8, 97-99].

By mapping ASH binding events during neural stem cell specification, we found a high co-occurrence with accessible regions pre-modelled during MZT, a time when Zelda is crucial for establishing chromatin organization for subsequent tissue-specific transcription [57, 100]. Since ASH proneurals are amongst the earliest zygotically transcribed genes [54, 101], we hypothesize that they may survey the early gastrula chromatin to gain access to neurogenesis related enhancers and possibly pre-initiate target transcription. This notion is supported by a single-cell RNAseq study of the early gastrula where the neuroectoderm primordium cell cluster expressed sc and some of its direct targets as identified here [102]. Later in the mature neuroectoderm, we demonstrate that proneurals also bind Zelda-independent elements, which showed restricted accessibility at the onset of zygotic transcription. ASH binding at these enhancers and concomitant gain in histone activation marks near known neural stem cell genes demonstrates their activating potency.

### ASC proneurals mediate the timely activation of the neural stem cell program in the neuroectoderm

Our work indicates that during NE to NB specification spanning stages 8-11, proneural-mediated chromatin reorganization and transcription is essential for the proper later unfolding of the entire NB lineage. For the first time we demonstrate that proneurals establish NB homeostasis of all 5 delamination waves, based on our genomic data (Fig. 2B), the phenotypic analysis of mutant NBs, both early (Fig. 4) and late born (Supplemental Fig. S6) and the expression patterns of the cloned proneural enhancers in vivo (Fig. 6). Thus, as reported for a single neuroblast, the MP2 [80, 81], it appears that all NBs that manage to delaminate in ASC mutants are mis-specified and cannot overcome functionally the initial stalling. Interestingly, murine Ascl1 depleted neural precursors also exhibit a similar delay [103]. Although proneural factors are crucial in the timely execution of the NB transcriptional program, partial activation of the program happens in their absence (Fig. 4). This is most likely mediated by different enhancers than those bound by ASH proteins (Fig. 6). The elusive proneural factors in ASC null embryos have been a long-standing puzzle [47, 104]. Such TFs could be Hb, in collaboration with Sna [105], since the expression of both was unaffected by ASC loss (Fig. 4A). Another possibility would be Daughterless, which heterodimerizes with ASH proteins, but also functions as a homodimer [52, 106]. Earlier observations have shown that L(1)sc and Ase can bind DNA as homodimers in vitro [107]. From the narrow overlap of our proneural binding consensus with Da (Supplemental Fig. S1G) it seems that in the embryonic neuroectoderm the two act to a large extent via distinct enhancers, contrary to the current belief that proneural factors are obligate heterodimeric partners of Da. This also agrees with the strong enhancement of the neural hypoplasia of double ASC and Da mutants [47]. On the other hand, it is unlikely that Wor and SoxN are the compensating proneural TFs as proposed by [104]. That study demonstrated that Wor and SoxN use their repressive capacities to promote neurogenesis, since EnR (Engrailed repression domain) fusions phenocopied their effect upon ectopic expression in epithelial cells [86]. It is unlikely that a duo of repressors would be able to activate the large cohort of NB specific genes that seems to be turned on by proneural factors (our study). In fact we have shown that *wor* is under ASH transcriptional control (Fig. 3C, Supplemental Fig. S3A) and reinstating its expression in ASC mutants is insufficient to rescue the CNS hypoplasia (Fig. 5C), although it mildly improves NB recovery. Regardless of the identity of other NB-promoting TFs, the eventual initiation of proliferation and rebound in the expression of key identity genes in ASC deficient NBs is insufficient to restore neural programming at the organism level, as evidenced by the depleted neuronal/glia progeny. This suggests that the ASC TFs are vital for neural stem cell homeostasis.

### Networks downstream of proneurals

Integration of the proneural binding events with the RNAseq and H3K27Ac changes during Notch mediated lateral inhibition revealed a downstream TF network, likely to consolidate the neural cell fate. Ase plays a central part in this network as being the only NB-specific TF with potent activating function [108]. The overlap of NE-expressed ASH binding events with NB-expressed Ase binding suggests that in the neuroectoderm ASH proneurals may mark neural enhancers which Ase will subsequently sustain to unfold the NB program. This is demonstrated in the sc19 deficiency where the presence of Ase partially improves mutant NB functionality and progeny development, compared to the deletion of all four ASC members (Supplemental Fig. S7). However, we find it impressive that the neuroectodermal ectopic induction of Ase can almost fully rescue the neurogenesis defects (Fig. 5F), proving, first, its functional equivalence to ASH TFs and, second, that the neural program must be installed early on during neural stem cell selection. The remaining TFs of this network are in their vast majority transcriptional repressors, highlighting the importance of blocking alternative transcriptional programs and differentiation fates to ensure proper unfolding of the NB program. We show that single members of this network contribute to neurogenesis, but we believe they mainly work combinatorically and in parallel to an ASC factor [86]. Snail TFs are central in this network and appear to have pivotal roles in NS development [70, 105, 109]. Snails however are not essential for NB ingression [67], instead, it seems that they regulate NB function and GMC transition [105, 110]. In addition to these core downstream TFs, NE proneurals bind near >1000 genes, which may contain previously uncharacterized players in implementing the NB fate and launching the subsequent GMC and neuron/glia developmental programs.

### Proneurals pioneer differentiation programs partly in the stem/progenitor cell

The mature VNC is the outcome of a complex crosstalk of glia and neuron signaling originating in the CNS, midline [75] and PNS [76]. Our identified proneural binding events near genes of all nervous sub-systems validate the genetic evidence of ASC involvement in their development [42, 47, 89, 111]. We thus propose that the late CNS defect in ASC embryos is the collective outcome of impaired stem cell specification and impaired progeny from different sub-systems, failing to establish the necessary communication cues.

In addition, studies in flies and mice have shown that, besides stemness, proneurals impact neuronal differentiation as well [112-116]. In our work, we identified binding near genes expressed in later differentiated cell types, GMC, neurons and glia (Fig. 2), where ASC gene expression has been extinguished. For at least one of these genes, *tap*, we showed that its protein expression is greatly compromised in ASC mutant GMCs (Supplemental Fig. S4C). We envision that this is happening in two ways: First, proneurals could regulate chromatin dynamics at neuronal/glial enhancers during neuroblast specification but robust transcriptional activation only happens later, delegated to TFs that appear as the neural differentiation program unfolds. Indeed, comparisons of chromatin states between stem cells and neurons support this notion. Some CNS-specific enhancers are “constitutive”, i.e. accessible from the NB all the way to neurons, whereas other neuron-specific enhancers gradually become accessible at later embryonic stages [5]. A second, not mutually exclusive, scenario is that key neuronal transcripts produced at the NB stage, are translationally repressed. Such genes are most likely pro-differentiation factors that generally lock cellular identity, as has been shown for the *elav* gene, whose transcription initiates in many cell types, but its protein product is strictly neuron-specific [117].

## CONCLUSIONS

We demonstrate that during stem cell specification ASC proneurals modulate chromatin dynamics to achieve the timely activation of neural transcription. This promotes stemness but also paves the way for appropriate lineage differentiation, which may explain the onset of developmental syndromes with ASCL1 mutations (OMIM: 209880). All stem cells and their future lineages within a tissue may depend on similar mechanisms of early chromatin remodeling, which is necessary for subsequent differentiation events.

## METHODS

### Drosophila stocks

UAS-CD8-GFP (II); bib-Gal4 (III) homozygous females were crossed to homozygous UAS-6xmyc-scAPAA, UAS-6xmyc-l(1)sc or UAS-NΔecd males for the embryo collections used in ChIPseq and RNAseq experiments. The Df(1)sc-B57 and Df(1)sc^19^ flies where rebalanced with a FM7,KrGal4,UAS-GFP chromosome to enable distinguishing the mutant embryos during imaging. Df(1)scB7/FM7,KrGal4,UAS-GFP(I); bib-Gal4(III) females were used for the UAS rescue experiments and for the UAS-FUCCI experiment.

For the generation of UAS-l(1)sc N-terminally 6xmyc-tagged flies, the l(1)sc coding region was amplified using primers with EcoR1 XhoI restriction sites overhangs (EcoR1-forward, XhoI-reverse) from yw cDNA (Superscript III, ThermoFisher 18080093), using KAPA High Fidelity Polymerase (Kapa/Roche, KK2103) and subsequently inserted in the entry pENTR™3C vector (ThermoFisher, A10464). We used pTMW (Drosophila Genomics Resource Center #1107) as the destination vector and the Gateway® LR Clonase® II kit (ThermoFisher, 11791020) to generate the final l(1)sc-pTMW vector. Subsequently the l(1)sc-pTMW construct was inserted into yw flies via P-element transformation . For the generation of enhancer-lacZ reporter flies we used the pBlueRabbit lacZ vector, which contains an hsp70 minimal promoter upstream of a lacZ reporter gene (Housden et al. 2012). Putative proneural bound regions were amplified with the corresponding primers with overhangs for EagI (forward primers) and XbaI (reverse primers) (see Table S7) from Oregon-R genomic DNA extracted with DNAzol™ (Theromofisher). PCR fragments were extracted from agarose gels (Macherey-Nagel, 740609.250). pBlueRabbit vector was digested with EagI and XbaI, gel extracted and dephosphorylated prior to ligations. Constructs were transformed using the φC31 integrase system into y w nos-int ; attP40[y+] / (CyO) hosts. All vectors generated for fly transgenesis were Sanger-sequence verified (Macrogen Inc). A complete list of fly strains and primer sequences are in Additional Supplemental Methods section.

### Embryo Collections, Immunostaining and Imaging

Embryo collections were made on cherry juice agar plates. Embryos were dechorionated in 50% bleach for 2 minutes. Dechorionated emrbyos were transferred to 4 ml glass tubes containing fixative solution (1200ul 1xPBS, 800ul 10% formaldehyde, 2ml heptane) and fixed for 20min with vigorous agitation. Embryos were devitellinized by vigorous shaking in methanol for 30-40secs. After 3 quick methanol rinses, samples were stored in methanol at - 20°C. On the day of immunostaining, embryos were rehydrated in PT (1xPBS, 0.2% Triton). Blocking was then conducted for at least 2 hours with PBT (PT+ 0.5% BSA). Primary antibodies were diluted in PBT and incubated overnight at 4°C. Next day, samples were washed extensively in PT. Embryos were incubated with secondary antibodies for 3 hours at room temperature. After extensive PT washes, 80μl n-propyl gallate-glycerol mountant was added to each sample and incubated overnight at 4°C. Embryos were then mounted and imaged in TCS SP8 confocal microscope system (Leica). Image analysis was performed with the Leica LAS X software. Antibodies used are listed in Additional Supplemental Methods section.

### ChIPseq protocol for low embryo number

We developed a low input Drosophila embryo ChIP-seq protocol based on [118]. Briefly, we set cages of 150 homozygous UAS-CD8-GFP (II); bib-Gal4 (III) female flies with 50 males homozygous for either UAS-scAPAA (II) or UAS-l(1)sc (II), or UAS-NΔecd (II), pre-conditioned for two days in vials before transfer to the cages. All embryo collections were performed during the same time window, from morning to mid-afternoon, to minimize clock-mediated changes in gene expression. A 30-minute preclearing step was performed every morning of collection. Egg lays were done on cherry juice/agar 6cm dishes for 0-3 hours at 27°C followed by a 3 hour maturation step at 29°C to boost GAL4 activity. We collected 3-6hs embryos on a Nitex mesh, dechorionated with 50% bleach for 2 minutes and washed with water. Subsequently, embryos were transferred with a brush in fixing solution and shaken for 10’ mildly in 2ml ependorfs. Fixing solution: 1500 μl Heptane, 100 ul 10% FA, 200 μl 10 X PBS and 200 double distilled H2O. Next, FA was quenched with glycine for 5’ minute with mild shaking. Fixing solution was discarded and embryos were washed twice with cold 1xPBS/0.1% Triton-X and then briefly low-speed centrifuged to pellet embryos. After discarding the second PBS wash, embryo pellets were stored in -80°C. A detailed protocol can be found in Additional Supplemental Methods section.

### Drosophila Embryo RNAseq

Embryos were collected at 0-2hs and then transferred to mature at 29°C for 3 hours (3-5hs collections). All embryo collections were performed during the same time window, from morning to mid-afternoon, to minimize clock-mediated changes in gene expression, after a 30-minute pre-clearing. Embryos were directly transferred in 50 μl Trizol containing tubes and stored at -80. On the day of RNA extraction, embryos were defrosted and homogenized using 1.5 ml manual pestle. For each replicate 5 independent daily collections were pooled after homogenization and RNA was isolated with phenol/chloroform without columns. RNA-seq libraries construction was performed with the Ion Total RNAseq Kit v2 (Thermo Fisher), using Poly(A) RNA selection with Dynabeads mRNA DIRECT Micro Kit Ambion (Life Technologies) according to manufacturers’ protocols. Libraries were sequenced on Ion Proton™ System (ThermoFisher) with PI CHIP v3, utilizing for template the Ion PI Hi-Q OT2 200 kit (# A26434) and the Ion PI Hi-Q Sequencing 200 kit (# A26433, A26772).

### NGS Data Analyses

Fastq files were transferred from Ion Proton to IMBB servers for storage and analysis. Mapping was performed to dm6 (UCSC/dm6, iGenomes, 2015). Software and Algorithms used in this study: SAMtools [119], MACS2 (v1.4) [120], HOMER (v4.5) [121], Hisat2 [122], Cutadapt (v1.12) (doi:https://doi.org/10.14806/ej.17.1.200.), HTSeq [123], edgeR [124], BEDTools [125], deepTools [126], GSEA (v4.0.3) [127], R (v4.0.3) (https://www.R-project.org/), Pavis (Flybase R6.01 assembly) [128], Flymine (v51) [63], i-cis Target [129], UCSC genome browser [130] (FlyBase/BDGP/Celera Genomics Release 6 + ISO1 MT), Flybase [64].

### ChIPseq Peak calling, Motif Analysis and Genomic Annotation

Mapping was performed using Hisat2 (--no-spliced-alignment --score-min L,0,-0.5), (samtools view -q 30). Bedgraphs were generated using bedtools genomecov and uploaded to the UCSC genome browser. Prior to peak calling, we excluded reads from the bam files mapped on repetitive regions. We also excluded reads that fell in our custom ‘black list regions’ (available upon request). Peak calling was performed using macs2 over input (-p 0.05) and peak overlaps were generated with bedtools (intersect -wa), excluding Chromosomes U and Uextra. The proneural consensus (Figure 1D) was generated imposing an FC>2 filter over input in the macs2 output file of the stronger second replicate of scAPAA. Motif analysis was done with homer findMotifsGenome.pl –size given. Assignment of peaks to genes was performed using homer annotatePeaks.pl. The genomic distribution of the datasets was performed by homer annotatePeaks.pl dm6 (default) and Pavis with parameters of upstream and downstream length set at 5 kb.

### Proneural Peak Consensus Overlapping with Zelda and chromatin marks during MZT

We overlapped our proneural binding consensus with Zelda binding events during blastoderm cellularization (the time of the maternal to zygotic transition) from two studies [54, 56] and found 41% and 62% overlap respectively. The proneural.vs.Zelda.Harrison data overlap was a subset of the proneural.vs.Zelda.Sun therefore we decided to continue with the second, presented in Figure 1, since it gave higher overlap with the proneural cistrome. We used the Table S5 from the Harrison study and the GSE65441_Zld_DESeq.txt.gz from the Sun study. Both datasets were converted to Drosophila genome version dm6 from dm3 using LiftOver in the UCSC browser.

### Proneural Consensus Overlaps with modENCODE datasets

For the DHS st5-st14 dataset [8] we downloaded the bed files of coordinates of 5% FDR peaks from UCSC/dm3 and then used LiftOver to convert to dm6. ChIP-seq data for Ac (ENCFF073ETO), Da (ENCFF718YZD), E(spl)m8 (ENCFF074INK) were downloaded from https://epic.gs.washington.edu/modERN/

### Heatmaps of ChIP datasets

We downloaded and mapped to dm6 parameters from the following Illumina sequencing datasets: SRR1779551 (Zelda) and its input SRR1779552. NC14 histone marks SRR1505729 (H3K27me3), SRR1505714 (H3K27Ac), SRR1505718 (H3K4me1) and SRR1505740 (input). SRR388356 (PolII) and SRR388382 (input). To correct for the difference in fragment size between Ion Torrent and Illumina sequencing we processed the IonTorrent datasets as follows: fastq reads were filtered and trimmed using cutadapt -m100 -l100 prior to Hisat2 mapping (--no-spliced-alignment --score-min L,0,-0.4 and samtools view -q 30). We indexed all bam files and used deepTools bamCompare, computeMatrix, plotProfile for Figure1H and S2E and plotHeatmap to generate Figures 1G and Figure 3B. We used as reference regions the center of proneural binding events (class I and II) ±5 kb from peak center. Heatmaps in Figures 1D and S1E were generated from the mapped reads, unprocessed for length, normalized over input, using ±5kb from proneural peak centers, using a custom script from the Odom lab [131], exported to images by TreeView software from the Eisen lab.

### Boxplot of ChIP datasets

For the boxplots in Fig. 3D, we used the multicov function of bedtools to count the processed trimmed reads from the H3K27Ac ChIP experiments on the stage 9 DHS dataset (Thomas et al, 2011). Subsequently, we generated the average read count per DHS from all 3 libraries and selected DHS sites that were in size equal to or greater than 50bp and had equal to or greater than 50 averaged reads per Kb of DHS. This filter resulted in 15,054 out of total 16,512 stage 9 DHS sites. Of these, 2,028 exhibited proneural binding (left), while 13,026 were not bound by proneurals (middle panel). Next we normalized the read counts within each DHS over the total number of uniquely mapped reads within library to correct for library size. 1,889 DHSs exhibited at least 1.3 fold change in U-scAPAA vs U-NΔE H3K27Ac ChIP datasets (right panel). Boxplots were generated using the log2 values of the corrected (for library size) reads counts in R (4.0.3). Statistics were performed with Wilcoxon rank sum tests.

### RNAseq Differential Analysis

Mapping was performed using Hisat2 (ref, --score-min L,0,-0.5). Counts were generated from bam files with HTSeq-count (-i gene_id). Differential Expression Analysis was performed with edgeR using batch correction and likelihood ratio tests (glmFit/glmLRT method), since replicates were performed in different time points resulting in large dispersions within groups. Tests were performed on 7,862 genes after keeping genes with cpm>3 in at least 3 samples. GSEA was performed on ranked gene lists from the edgeR output files using BDGP gene ids and genes assigned to proneural peaks or affected DHS sites.

## DATA ACCESS

The sequencing data generated in this study have been submitted to the NCBI BioProject database (https://www.ncbi.nlm.nih.gov/bioproject/) under accession number PRJNA719934.

## ACKNOWLEDGEMENTS

We thank Ioannis Livadaras for embryo injections and Maria Monastirioti for critical reading of the manuscript. Manolis Dialynas for data management. We thank the following students that contributed to experimental procedures: Efstathia Mpampoula, Eva Ioannou, Konstantina Mylonaki, Konstantinos Klaourakis, Krystallia Gourlia, Mary Chatzi, Christina Kosmopoulou, Christos Zioutis, Florentia Romanou and Christina Thomou. We thank Margarita Stapounzi for technical assistance. We thank Eirini Stratidaki and Niki Gounalaki at the IMBB Genomics Facility for library preparation and sequencing. We also thank Pantelis Hatzis from the Genomics Facility of Alexander Fleming Research Center Institute in Athens for library construction and sequencing one RNAseq replicate. Alexander Babaratsas for Drosophila stock maintenance.

## AUTHOR CONTRIBUTIONS

V.T. designed, supervised, performed and analyzed the majority of experiments, wrote and prepared the manuscript. K.S., M.D., D.T. performed experiments. C.D. designed, performed experiments, analysed data and wrote the manuscript.

