## Supplemental Figures for "ASC proneural transcription factors mediate the timely initiation of the neural program during neuroectodermal to neuroblast transition ensuring progeny fidelity"

**Supplemental Figures and legends (Fig. S1-S8)**

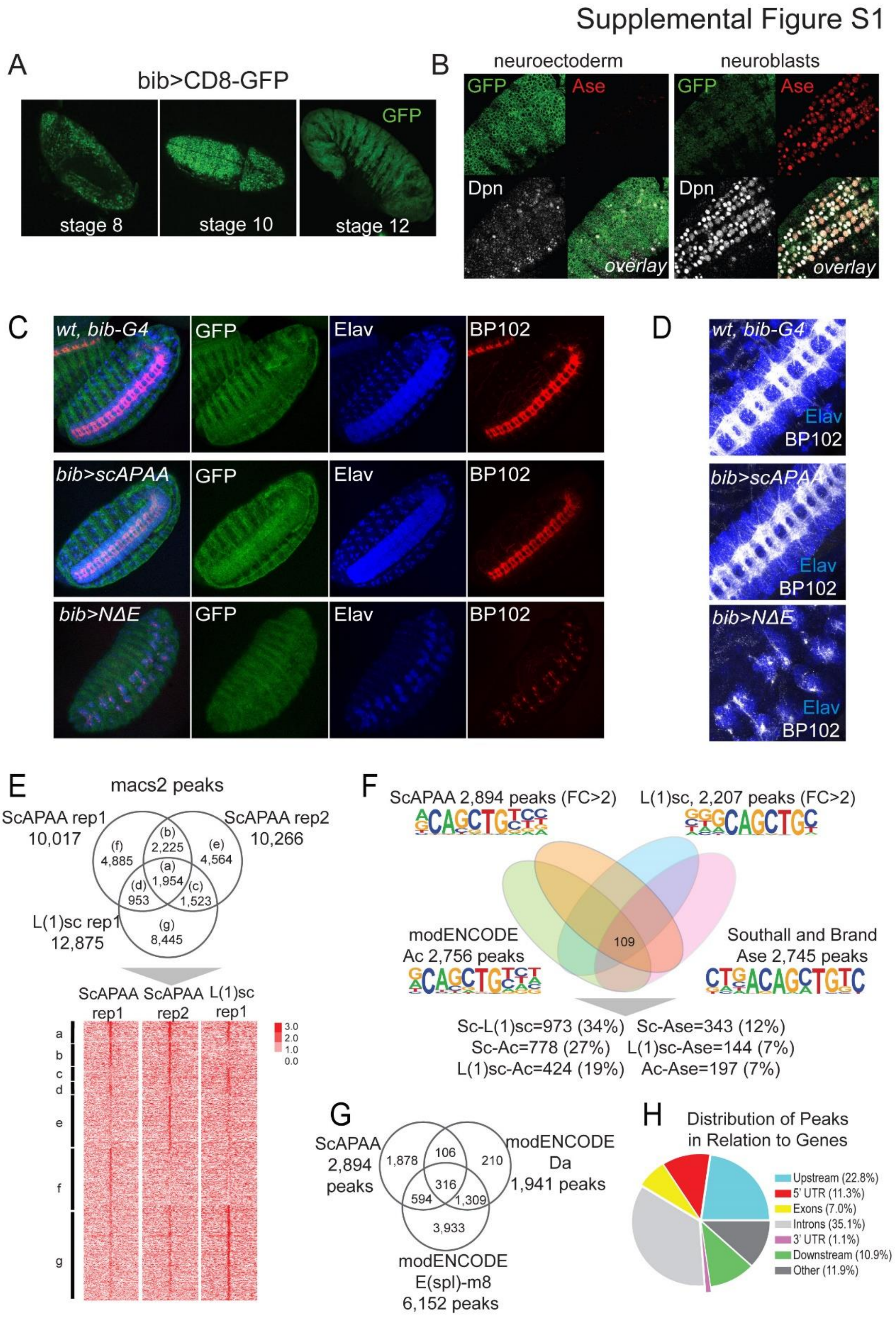


**Fig. S1. Phenotypes of bib-Gal4 driven expression of U-scAPAA and U-NΔE and genomic analyses of proneural binding consensus.** A) Stage 8, 10 and 12 embryos of bib-GAL4 x U-CD8-GFP show neuroectodermal expression, which expands laterally at late stages (st 12). B) Stage 10 neuroblasts in bib-GAL4 x U-CD8-GFP express residual GFP. Single sections, using the same laser settings, for the neuroectoderm and neuroblast focal planes of the same embryo. C) Stage 16 embryos showing mild nerve cord hyperplasia in U-scAPAA and severe hypoplastic phenotype in U-NΔE;Elav marks neuronal nuclei; BP102 marks axons. D) Neuromeres of stage 16 embryos of U-NΔE embryos exhibit clumps of neurons with limited axonogenesis compared to wt or U-scAPAA. E) Peak overlaps among the 3 replicates of ScAPAA and L(1)sc ChIP experiments. Heatmap below the venn diagram shows the corresponding normalized over input signals of read density. F) Overlap at the level of peaks between all 4 ASC family members and the percentages of pairwise comparisons – percentages state the overlap of binding events for the first factor with respect to the second. De novo motif analysis (homer) revealed slight changes in the enriched E-box bHLH motif. G) Venn diagram of binding peaks overlaps among the proneural ScAPAA consensus and Da and E(spl)m8 from modENCODE data. H) Genomic distribution of the 2,984 proneural binding consensus (Pavis).

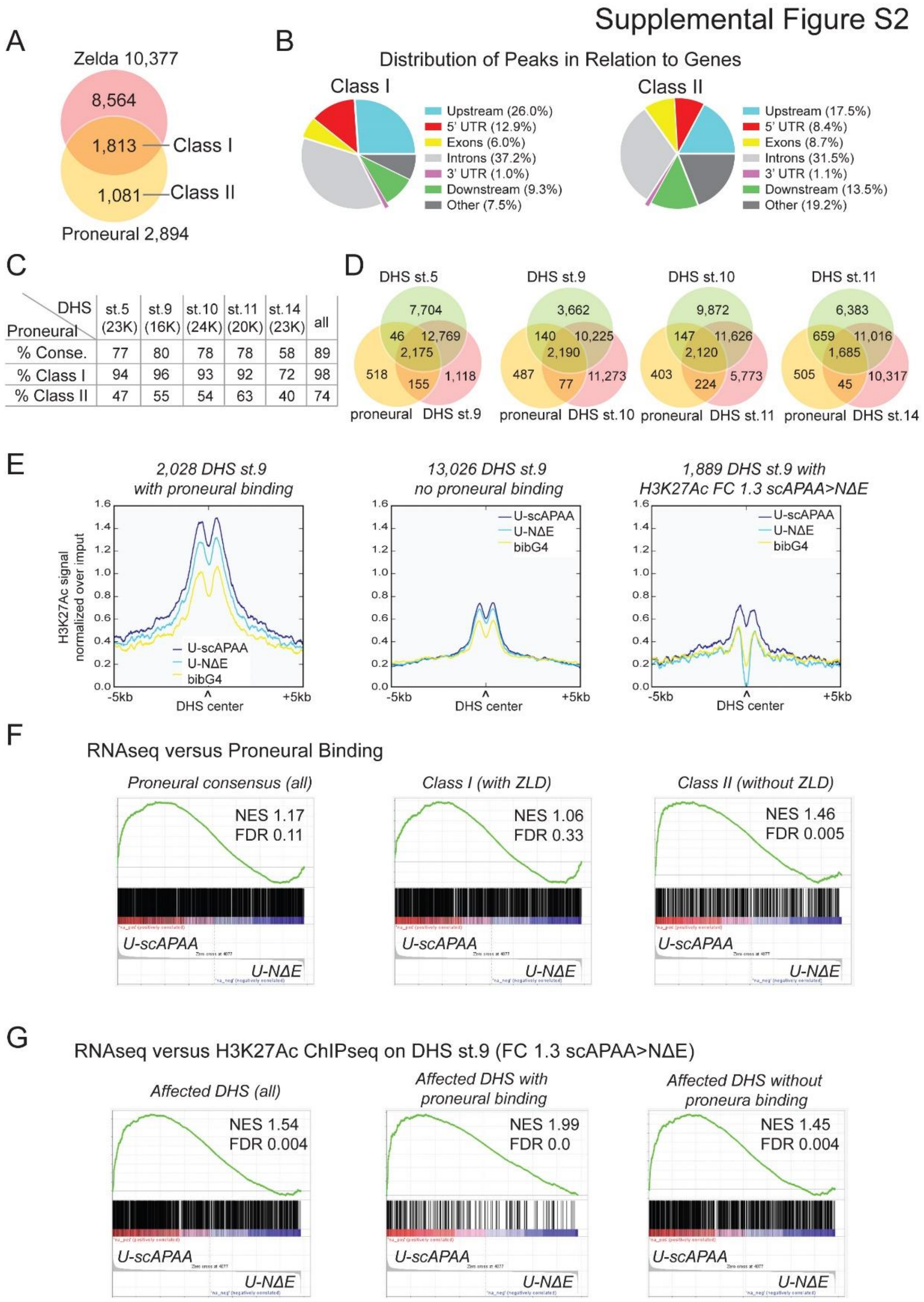


**<u>Fig. S2</u>. Proneural regulated chromatin effects correlate with transcriptional output**
A) Overlap of proneural consensus peaks during neuroblast specification with Zelda binding events during MZT. Two classes of events are shown, Class I bound by Zelda and class II Zelda independent. B) Genomic distribution of class I and II proneural binding sites (Pavis). C) Percentage of proneural binding peaks overlapping with stage specific DHS sites from Thomas et al (2011), The top row refers to all 2,894 peaks; the next two rows present Class I and Class II peaks separately.  D) Venn diagrams of proneural peaks with consecutive stage

specific DHS sites. E) Averaged, normalized signal of the H3K27Ac ChIPseq from U-scAPAA, U-NΔE and wt, bibG4 embryos in stage 9 DHSs: 2,028 with proneural binding (left), 13,026 without proneural binding (middle) and in 1,889 DHS loci that exhibited FC 1.3 in U-scAPAA>U-NDE. F) Gene set enrichment analysis (GSEA) plots of the RNAseq ranked genes in U-scAPAA vs. U-NΔE with all Proneural binding events (left), class I (middle) and class II (right). G) GSEA plots of the RNAseq ranked genes in U-scAPAA vs. U-NΔE with stage 9 DHS sites affected in H3K27Ac by FC>1.3 in U-scAPAA>U-NDE, all (left), 306 that are also proneural bound (middle) and 1,583 affected DHS sites but not bound (right).

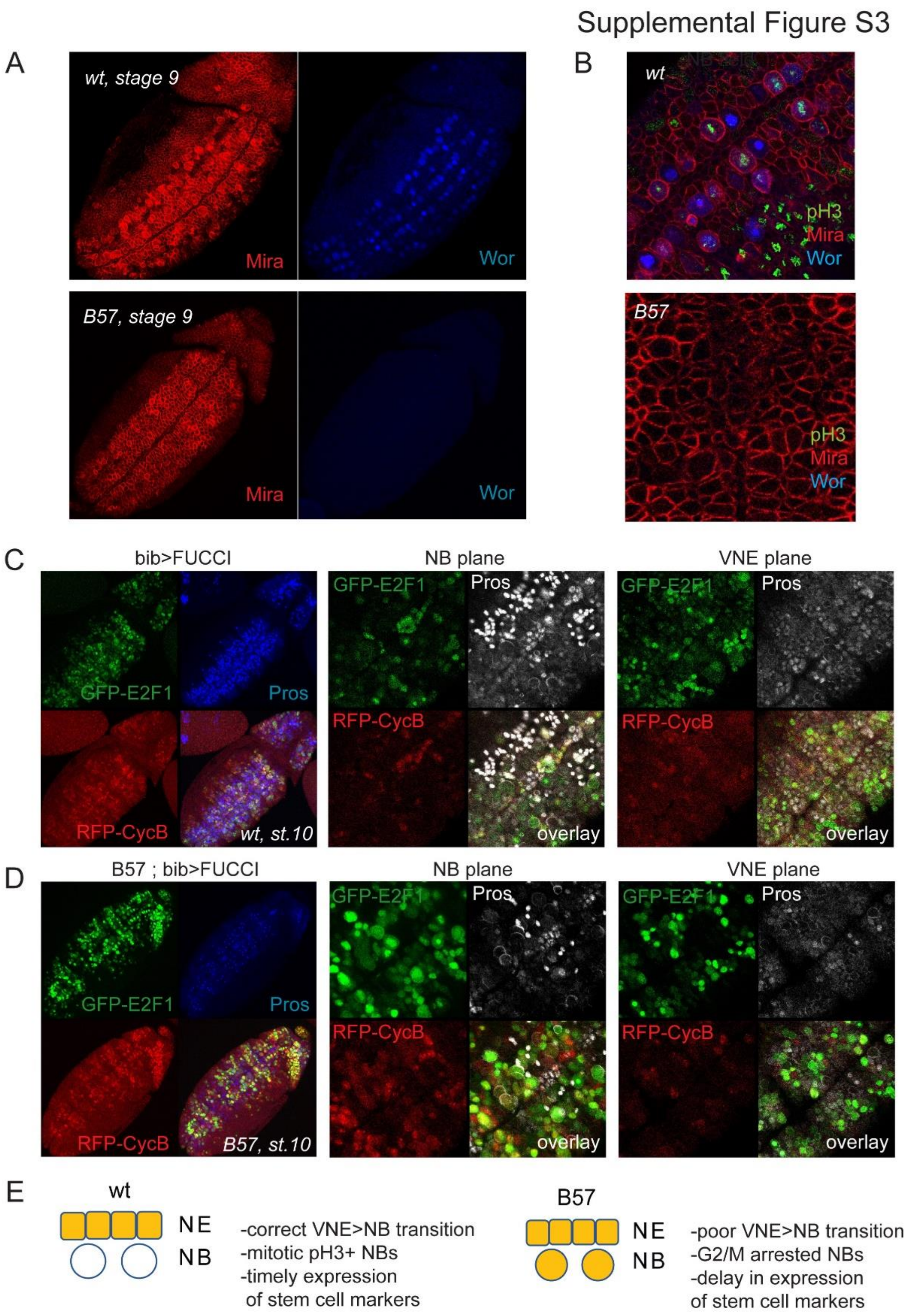


**<u>Fig. S3</u>. ASC mutant neuroblasts are initially arrested at G2/M.**
Stage 9 S1 wt neuroblasts express Worniu robustly, whereas mutants do not. Mira stains the entire ventral NE. B) Single sections at the neuroblast level below the embryo surface shows that S1 delaminated neuroblasts of Df(1)scB7 embryos do not express Worniu nor do they proliferate (pH3), unlike wt neuroblasts. C) bibGal4 wt stage 10 embryo expressing a dual UAS-GFP-E2F1;UAS-RFP-CycB (FUCCI). Left panel shows a projection of a ventral view of the entire embryo, where GFP and RFP are expressed in the neuroectoderm in response to bibGal4. At the NB level (middle panel) limited GFP/RFP expression is detected. Right panel

shows strong signal of GFP (G1 marker) at the overlying ventral neuroectoderm level. D) In Df(1)scB7 stage 10 embryo (left) which is just rebounding (few Pros+ GMCs present), many delaminated neuroblasts (middle) that have not divided yet co-express both GFP and RFP suggesting a G2/M arrest, whereas the VNE plane (right) is comparable to wt. E) A model cartoon summarizing the phenotype of the delaminated cells in the deletion of ASC proneural genes.

Supplemental Figure S4

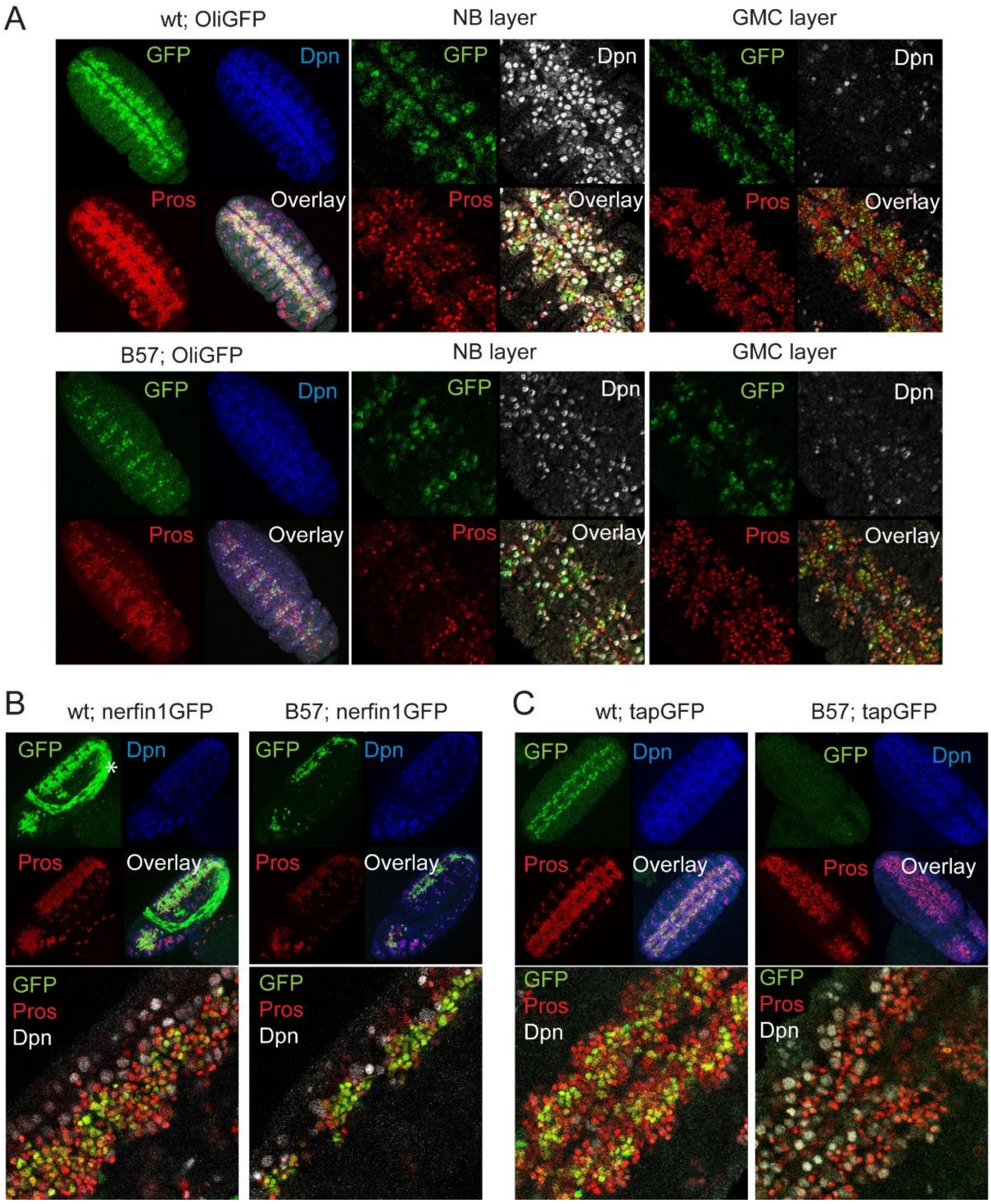


**<u>Fig. S4</u>. GMC expression of proneural targets is also impaired in ASC mutants**

A) OliGFP exhibits neuroblast and GMC expression in wt embryos. In Dfsc(1)B57 after the stalling window OliGFP expression is restricted. B) nerfin1GFP is predominantly expressed in the GMC pool and shows restricted expression in Dfsc(1)B57 embryos. The GFP in the amnioserosa (marked with *) is coming from the FM7, KrGAL4,UASGFP chromosome used to distinguish wt from mutant embryos. C) tapGFP is normally expressed in many GMCs and in Dfsc(1)B57 embryos its expression is not observed during stage 11.

Supplemental Figure S5

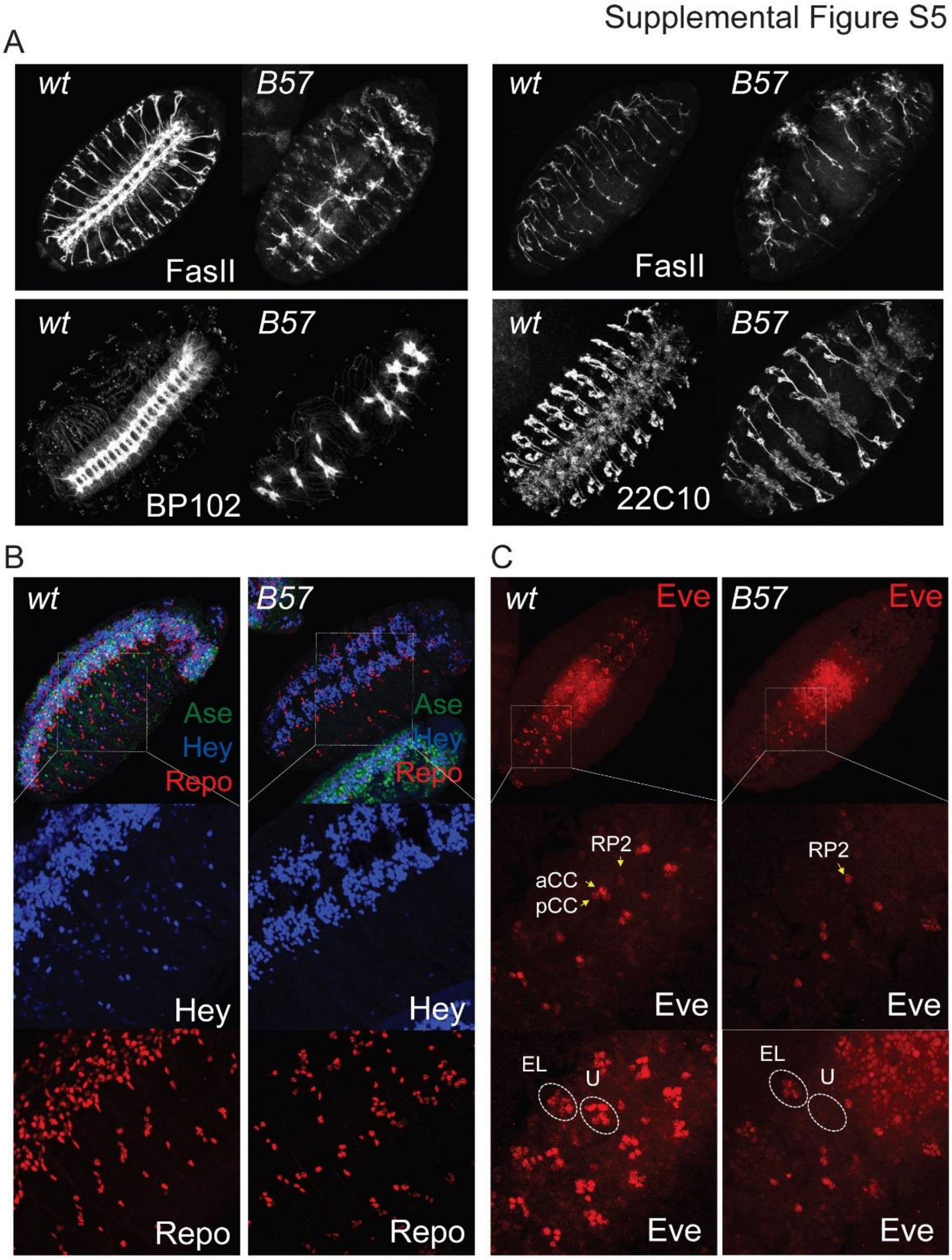


**<u>Fig. S5</u>. Depletions of the glia and neuronal populations in ASC mutants result in severe axonal defects.** A) Stage 16 Dfsc(1)B57 embryos, stained with FasII, BP102 and Futsch/22C10 display extensive neuronal hypoplasia. Note the complete lack of the longitudinal VNC tracts (FasII and BP102 ventral views, left panels), the defects in intersegmental and the segmental nerve development and pathfinding (FasII sagittal, top right panels) and the tightly apposed bilateral neuronal populations in the VNC neuromeres suggestive of incomplete midline function and production of repulsion cues (22C10 ventral view, bottom right panels). B) Repo (glia) and Hey (neuron subset) staining reveal a severe loss in differentiated cell populations. C) The Eve positive neurons are diminished in Dfsc(1)B57 embryos. Middle panels are closeups of the dorsal VNC; bottom panels ventral VNC of the same field.

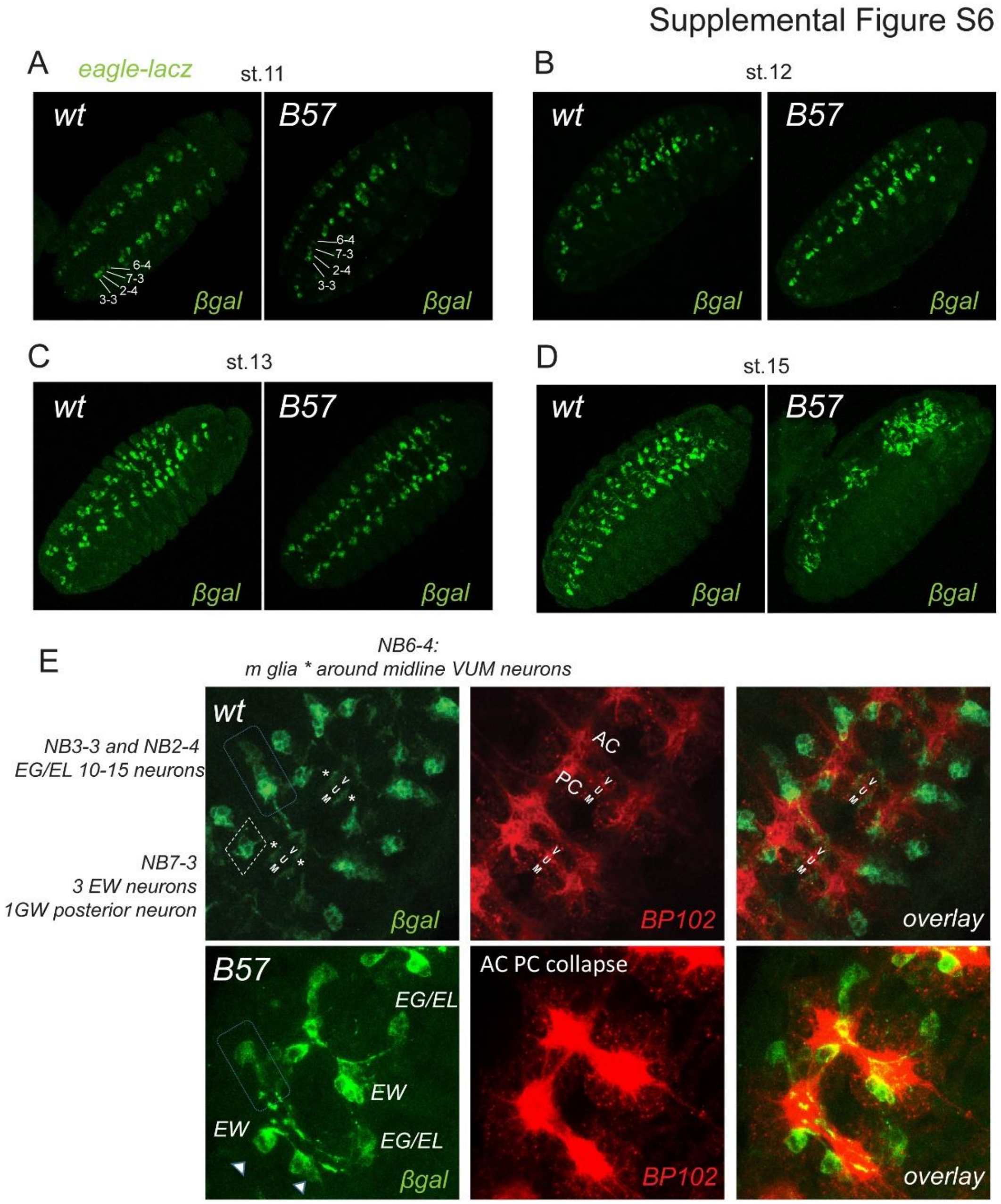


**<u>Fig. S6</u>. Proneurals contribute to late NB identity and progeny fidelity**

A) eagle-lacZ expression shows that at stage 11 all four positive neuroblasts have delaminated in Df(1)scB57. B) Stage 12 embryos do not show major differences in eagle-lacz positive cells. C) By stage 13, the medial glia, progeny of 6-4, that move towards the midline are absent or have lower expression in the Df(1)scB57. D) By stage 15 the nerve cord shows severe disorganization in mutants. E) Stage 15 VNC close-ups show an Anterior-Posterior Commissural axonal collapse, no medial glia (asterisks in wt), a diminished EG/EL neuronal population (boxed), an EW neuron missing (left arrowhead) whereas the other EW (right arrowhead) is sending its axon laterally instead of posteriorly.

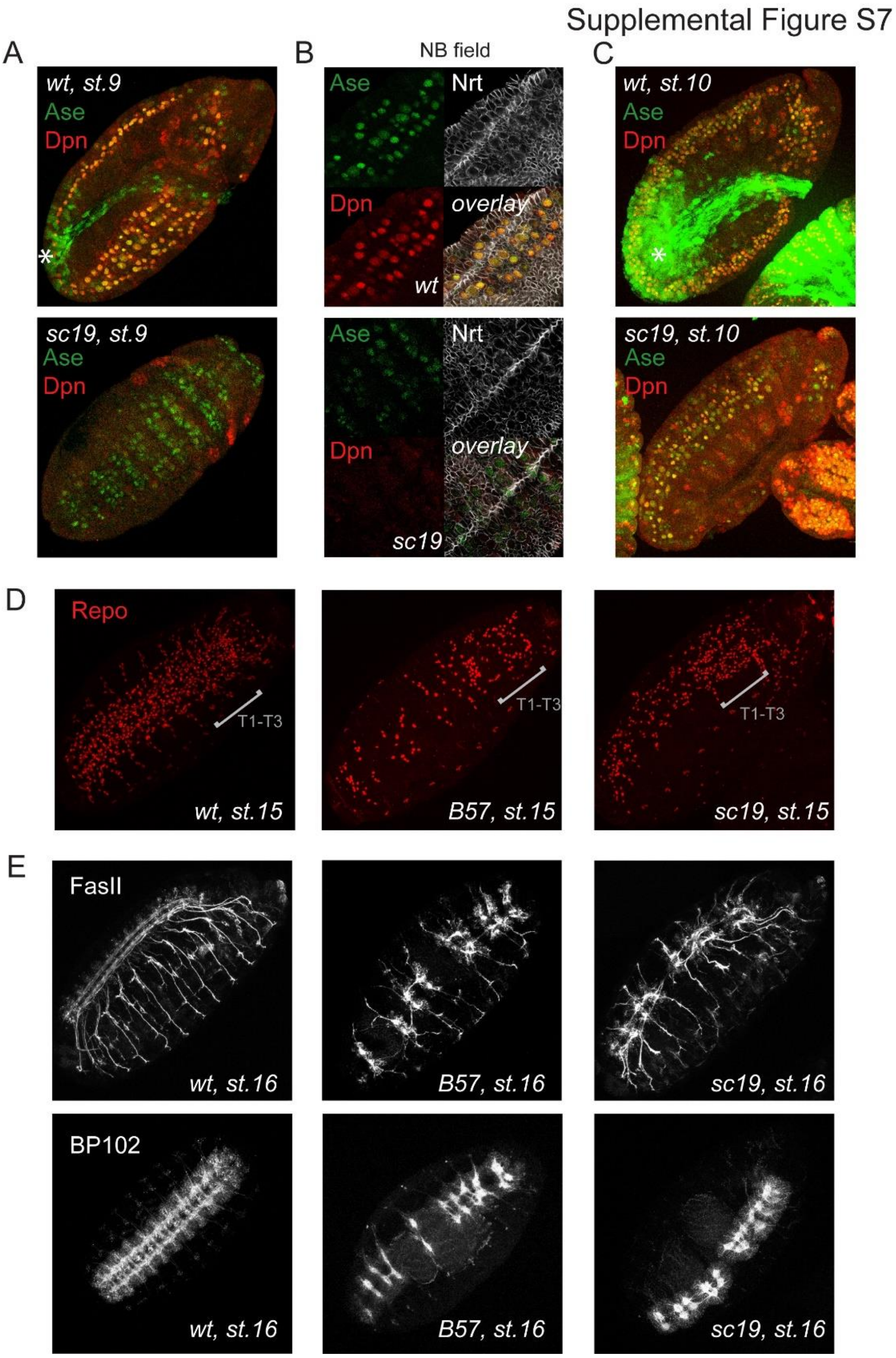


**<u>Fig. S7</u>. Asense provides partial neuroblast functionality and CNS development in the absence of ac, sc and l(1)sc.** A) Stage 9 wt and Df(1)sc19 embryos. Wt neuroblasts express Ase and Dpn robustly while mutant have weak transient expression of Ase at the NE layer. B) Single sections at the neuroblast level show that in Df(1)sc19 embryos not all neuroblasts (large round cells) express Ase and none express Dpn. Nrt marks cell outlines. C) During stage 10 both wt and Df(1)sc19 embryos express both Dpn and Ase in neuroblasts. Note that the wt embryo also shows KrGal4>GFP expression (used to distinguish wt from mutant embryos) in the green channel, noted with a *. C) Df(1)sc19 embryos have more glia than Df(1)scB57 E) Axonal hypoplasia is less severe in Df(1)sc19 compare to Df(1)scB57.

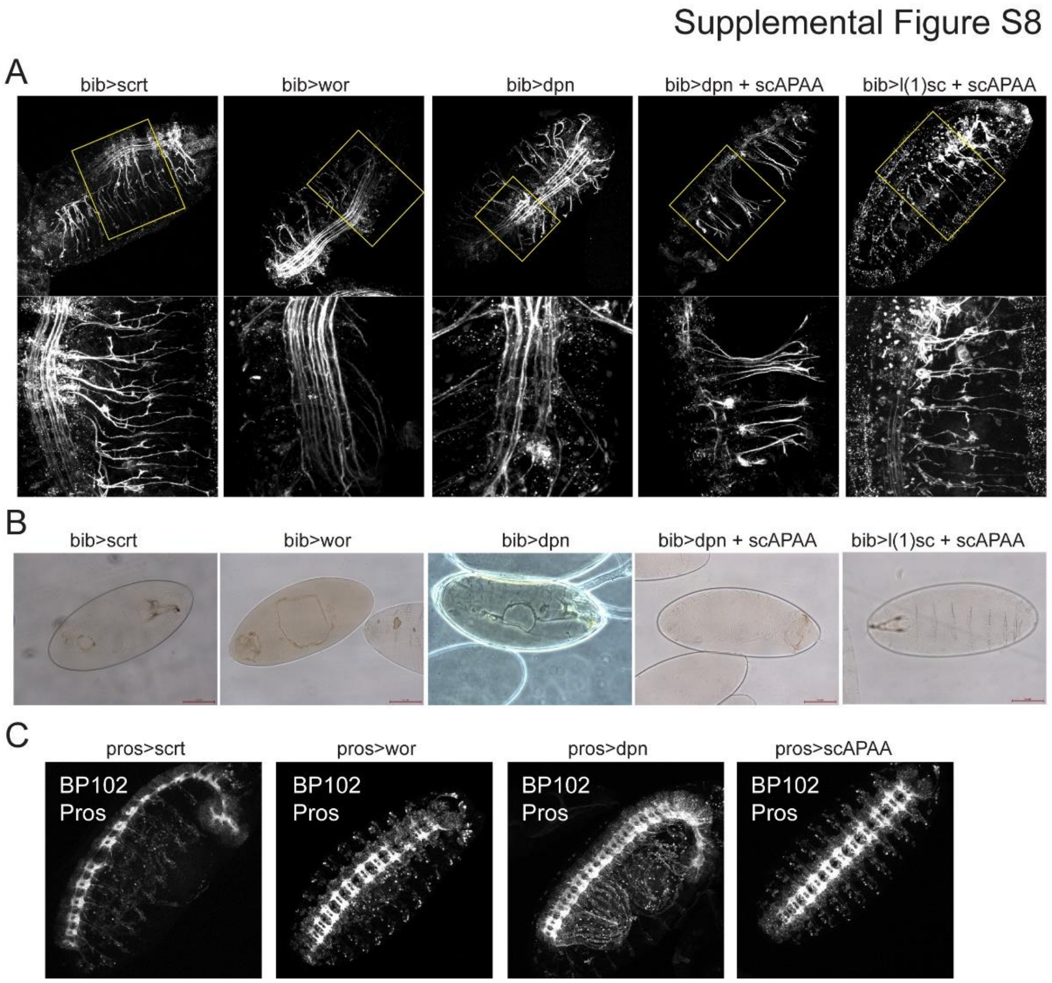


**<u>Fig. S8</u>. Neuroectodermal induction of proneural targets enhances neurogenesis at the expense of epidermal fate.** A) Neuroectodermal overexpression using bibGal4 of UAS-scrt, UAS-wor, UAS-dpn, UAS-dpn + UAS-scAPAA and UAS-l(1)sc + UAS-scAPAA result in highly hyperplastic late CNS phenotypes. A) Cuticle preparation from bibGal4 UAS-scrt, UAS-wor, UAS-dpn and UAS-dpn + UAS-scAPAA show ventral/ cephalic holes indicative of defects in the epidermis. C) Induction of UAS targets in the neuroblasts using pros-Gal4 does not affect late CNS development.
